# CD28 Co-stimulation Is Organized by Signaling Condensates and Counteracted by PD-1

**DOI:** 10.64898/2026.02.03.703677

**Authors:** Hui Chen, Songfang Wu, Yong Zhang, Wei Hu, Changjie Lou, Wei Chen, Jizhong Lou

## Abstract

T cell activation relies on the coordinated integration of TCR and co-signaling pathways, yet how these signals are physically organized remains unclear. Here, we demonstrate that CD28 co-stimulation is driven by a phase-separation mechanism in which the CD28 intracellular domain (ICD) forms signaling condensates with the Src-family kinase Lck. Ligand engagement promotes the CD28 ICD accessibility, and triggers rapid CD28/Lck condensation, enhancing CD28 phosphorylation, cytoskeletal polarization, and cytokine production. We further identify PD-1 as a graded regulator of this process: monophosphorylated PD-1 is recruited into CD28/Lck condensates, whereas di-phosphorylated PD-1 competitively binds the CD28 ICD and dissolves these assemblies. CD28 and PD-1 ICDs also undergo condensation directly, revealing a dynamic and compositionally heterogeneous signaling system. Guided by this competitive phase-separation mechanism, we engineer CD28 ICD mutants that preserve Lck condensation while resisting PD-1-mediated disruption. Incorporation of these condensation-selective mutants into CAR constructs confers resistance to inhibitory signaling and enhances antitumor efficacy in vivo. These findings establish phase separation as a central organizing principle of CD28 co-stimulation and provide a biophysical framework for engineering more resilient therapeutic T cells.

## Introduction

T cell activation is initiated by the engagement of peptide-MHC complexes with the T cell receptor (TCR) and is subsequently shaped by a variety of regulatory signals, including co-stimulatory or co-inhibitory receptors, cytokines, and more recently discovered metabolic cues (Chen and Flies, 2013; Shi et al., 2024). Harnessing this activation machinery has enabled the development of adoptive T cell therapies, which have achieved notable clinical success in cancer and autoimmune diseases and now represent one of the most promising immunotherapeutic modalities (Rosenberg et al., 1988). Advances in T cell engineering have further expanded the capacity of T cells to recognize and eliminate target cells (Eshhar et al., 1993; Sadelain et al., 2017). Three major classes of engineered T cells — CAR-T cells, TCR-T cells and STAR-T/HIT-T cells — have demonstrated substantial therapeutic benefits in oncology (Finck et al., 2022; Liu et al., 2021). Notably, second-generation CAR-T cells incorporating co-stimulatory signaling domains such as CD28, 4-1BB, or their combinations thereof have markedly improved T cell persistence and cytotoxicity (Imai et al., 2004; Maher et al., 2002; Milone et al., 2009). More recently, analogous co-signaling modules have been introduced into TCR-T, TRuC-T, and STAR-T/HIT-T platforms, underscoring the central role of co-stimulatory signaling in engineered T cell function (Dobrin et al., 2024; McCarthy et al., 2023; Mog et al., 2024).

CD28 was first identified in 1980 by Hansen as a T cell-restricted surface antigen and was later recognized as a crucial co-stimulatory receptor required for optimal T cell activation and proliferation (Weiss et al., 1986). Upon engagement of its ligands (CD80 or CD86) either in-*trans* or in-*cis*, the intracellular domain (ICD) of CD28 undergoes release from the plasma membrane, exposing the YxxM and PYAPP motifs for phosphorylation by the Src-family kinases Lck and Fyn (Dobbins et al., 2016; Yang et al., 2017; Zhao et al., 2019). Phosphorylated CD28-ICD then recruits PI3K, Grb2 and Gads to initiate downstream signaling cascades (Azuma et al., 1992; Boomer and Green, 2010; Raab et al., 1995; Sadra et al., 2004; Zhao et al., 2023). CD28 signaling is tightly constrained by multiple regulatory pathways: the phosphatase CD45 attenuates CD28 signaling and limits T cell proliferation (Turka et al., 1992; Vandenberghe et al., 1992); CTLA-4 competes with CD28 for CD80/CD86 binding, thereby preventing CD28 activation (Schwartz et al., 2001; Stamper et al., 2001); and the inhibitory receptor PD-1 recruits SHP-2 to dephosphorylate the CD28-ICD, resulting in potent suppression of CD28-dependent signaling (Hui et al., 2017a).

T cell activation is spatially organized within the immune synapse (IS), a highly structured “bulls-eye” interface formed between T cells and antigen-presenting cells, characterized by a central accumulation of TCR signaling components at the center, surrounded by co-signaling receptors and adhesion molecules (Dustin and Depoil, 2011). We and others have shown that key TCR-proximal signaling events are governed by biomolecular condensation and phase separation. We previously described a self-programed condensation of TCR/Lck complexes that enhances TCR clustering and phosphorylation (Chen et al., 2023). Following TCR engagement, phosphorylated CD3 chains recruit Zap70, which phosphorylates the adaptor molecule LAT. Phosphorylated LAT then acts as a scaffold to promote phase separation with PLCγ1, GRB2, and SOS1, driving downstream T cell activation (Su et al., 2016). In parallel, phosphorylated CD3ε recruits Csk, providing a negative feedback mechanism that restrains Lck activity and prevents excessive signaling (Chen et al., 2023). Co-inhibitory receptors also exploit phase-separation mechanisms: Lag-3 undergoes condensation with CD3ε to disrupt TCR/Lck condensates (Du et al., 2025). Additionally, the co-receptors CD4 and CD8 have been reported to undergo phase separation with Lag-3, leading to their disassociation from Lck, thereby impairing TCR signaling and T cell activation (Guy et al., 2022). More recently, TIGIT was reported to form CD155-dependent condensates that coalesce with the TCR during T cell activation (Worboys et al., 2023). Whether classical co-stimulatory receptors similarly employ phase-separation-based mechanisms to regulate T cell activation remains unclear.

Here, we demonstrate that the co-stimulatory receptor CD28 undergoes phase separation with Lck to form signaling condensates that promote T cell activation. We further show that phosphorylated PD-1 negatively regulates CD28/Lck condensation through a ligand-independent, competitive interaction with the CD28 intracellular domain. Disrupting this CD28–PD-1 interaction via targeted mutations within the CD28-ICD enhances T cell resistance to PD-1-mediated inhibition. Moreover, incorporation of this CD28 mutant into a second-generation CAR-T construct results in increased cytokine production and improved therapeutic efficacy. Together, these findings identify a previously unrecognized condensate-based mechanism for CD28 signaling regulation and provide a rational strategy to enhance the performance of engineered T cell therapies.

## Results

### CD28 forms condensates with Lck in-vitro and in cells

We previously established a supported lipid bilayer (SLB) system that enables immobilization of Lck kinase (Lck_UD-SH3-SH2_, residues 3-226 or full-length Lck-wt, residues 3-509) (Chen et al., 2023), allowing us to evaluate whether the ICDs of membrane receptors undergo phase separation with Lck (Figure S1A). To determine whether co-signaling receptors can condense with Lck, we examined the ICDs of six representative co-signaling receptors, three co-stimulatory (CD28 residues 180-220, 4-1BB residues 214-255, ICOS residues162-199) and three co-inhibitory (CTLA-4 residues 183-223, PD-1 residues 192-288 and Lag-3 residues 472-525). Among all ICDs tested, CD28-ICD robustly phase-separated with both both Lck_UD-SH3-SH2_ and Lck-wt on SLB (Figure 1A-1C).

**Figure 1.**
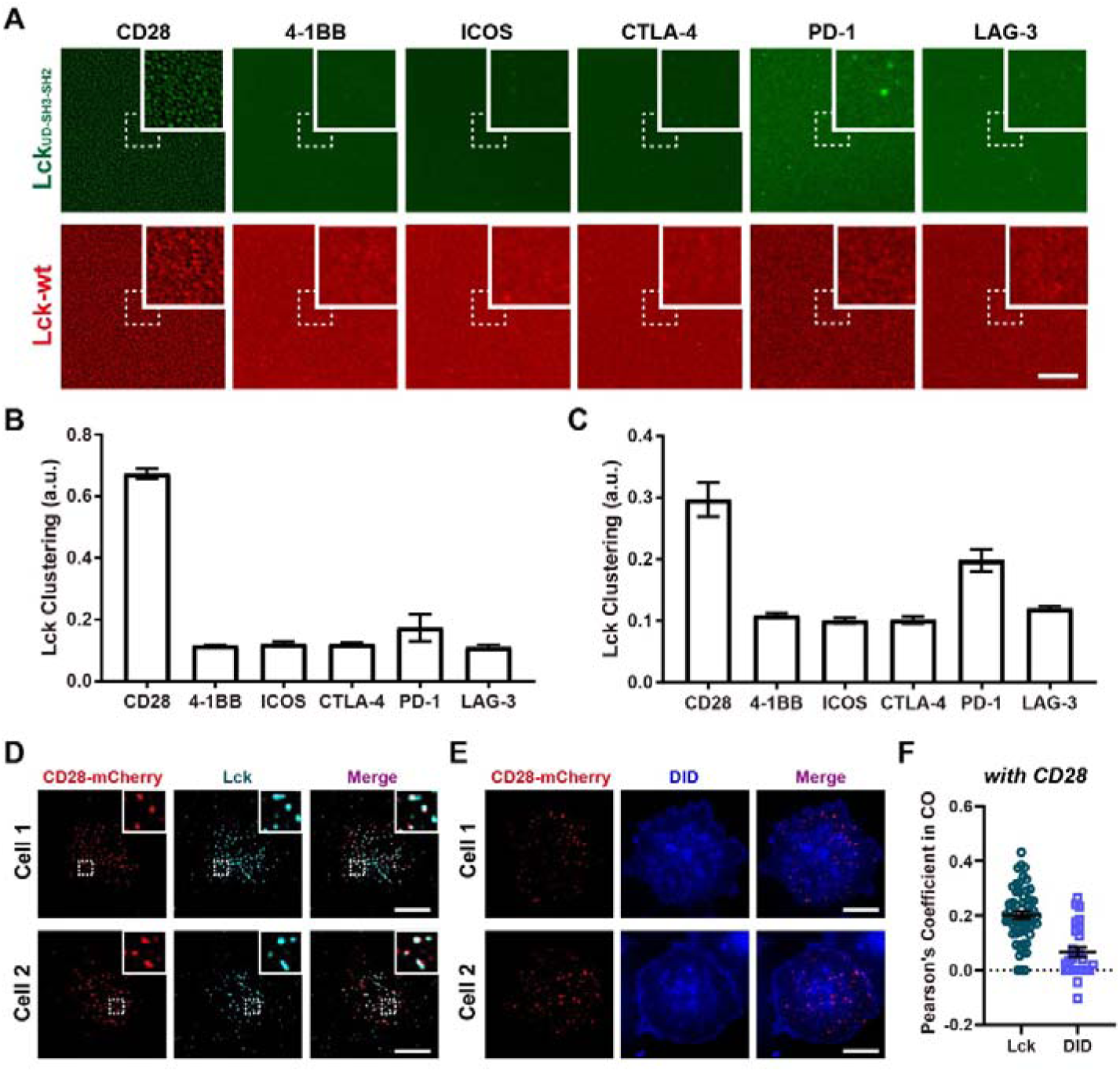
CD28 undergoes phase separation with Lck. **A-C**: Representative images (**A**) and quantification (**B**-**C**, n=5) of phase separation between the ICDs of co-signaling receptors and Lck_UD-SH3-SH2_ (**A**, upper panel; **B**) or Lck-wt (**A**, lower panel; **C**) on SLBs. Scale bar: 10 μm. Data are presented as Mean±SD. **D-E**: Representative images of Jurkat-CD28^mCherry^ cells stimulated on anti-CD28 coated glass surface and stained with anti-Lck (**D**) or the membrane dye DID (**E**). The co-localization of CD28 with Lck (n=55) and membrane dye (DID, n=26) were quantified by calculating Pearson’s coefficient (**F**). Scale Bar: 10 μm, data are presented as scatter plot with Mean±SEM.

To further confirm CD28/Lck colocalization, we next immobilized His-tagged CD28-ICD on SLBs. FRAP analysis showed that CD28 immobilization did not alter SLB dynamics (Figure S1A and S1B). Lck_UD-SH3-SH2_ strongly enriched within CD28-ICD clusters, supporting CD28/Lck phase separation (Figure S1C).

We next examined whether CD28/Lck condensates form in cells. Full-length mCherry-tagged CD28 was over-expressed in CD28-Knockout Jurkat cells (Jurkat-CD28^mCherry^) and stimulated on glass surfaces coated with anti-CD28 antibody. CD28 and Lck formed clear co-clusters following stimulation (Figure 1D). Membrane staining indicated that these puncta were not caused by membrane invaginations (Figure 1E and 1F).

### The BRS motif of CD28-ICD regulates CD28/Lck clustering and CD28 signaling

The CD28-ICD contains several conserved motifs, including YxxM and PYAPP motifs required for downstream signaling; two basic-rich sequences (BRS), BRS1 and BRS2, and proline-rich sequence (PRS) that interacts with Lck or Fyn to regulate CD28 phosphorylation (Figure S2A and S2B). To define motifs required for phase separation, we generated a panel of CD28-ICD mutants and tested their ability to condense with three Lck truncations (Lck_UD_, Lck_UD-SH3_ and Lck_UD-SH3-SH2_). BRS1 proved essential for CD28/Lck clustering, wheras BRS2 contributed primarily to clustering with Lck_UD_ (Figure 2A and 2B).

**Figure 2.**
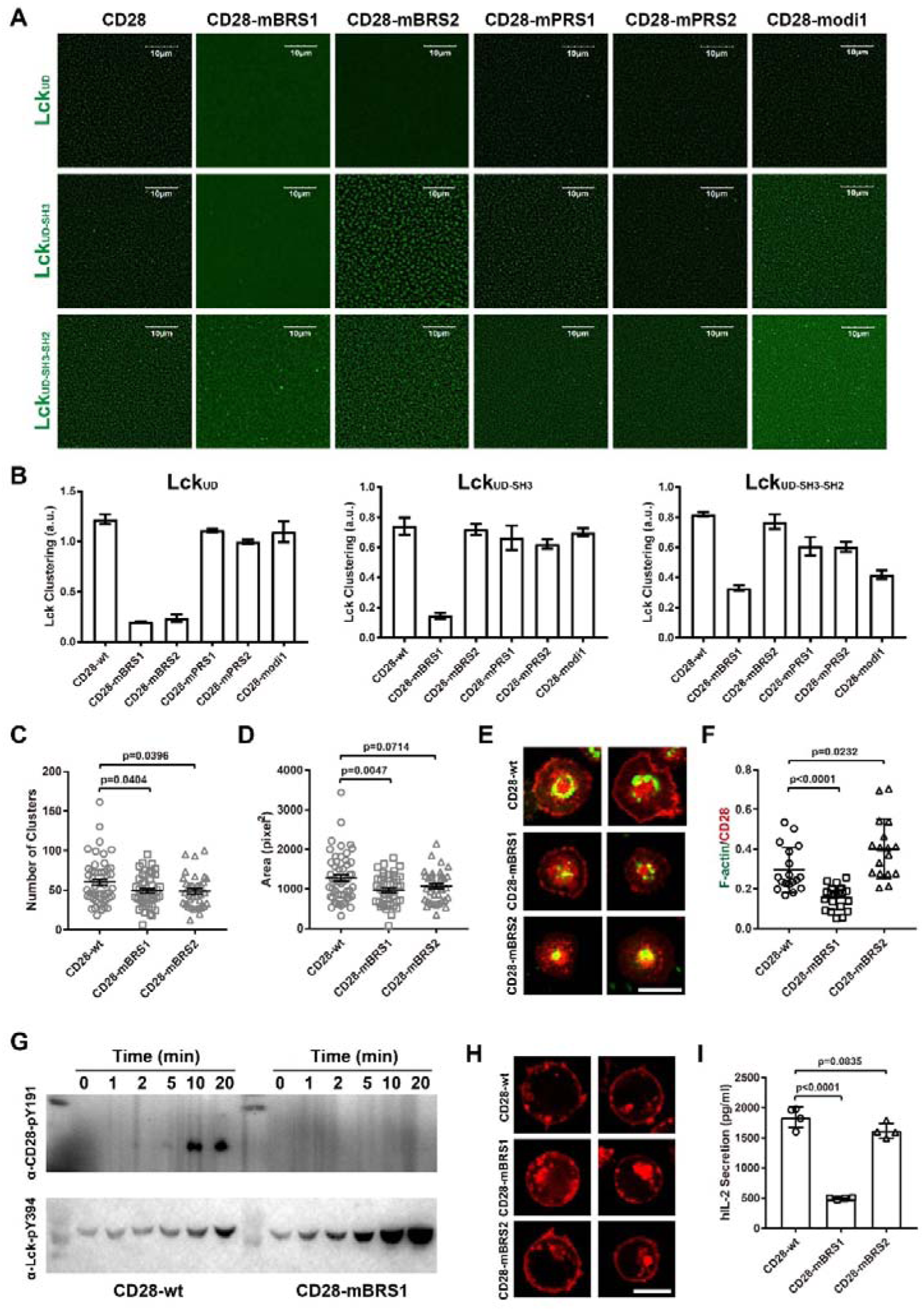
BRS of CD28 controls CD28/Lck condensation and downstream signaling. **A**-**B**: Representative images (**A**) and quantification (**B**, n=5) of condensates formed by CD28-ICD mutants with Lck truncates (AF488 labeled). Scale bar: 10 μm. Data are presented as Mean±SD. **C**-**D**: Quantification of CD28 clustering in Jurkat-CD28^mCherry^ cells stimulated on anti-CD28 coated glass. Shown are the number of CD28 clusters (**C**) and cluster area (**D**). CD28-wt: n=52; CD28-mBRS1: n=44; CD28-mBRS2: n= 37. Data are presented as Mean±SEM. **E**-**F**: Representative images (**E**) and quantification (**F,** n=18, 20, 17) of actin polarization in Jurkat-CD28^mCherry^ cells stimulated on anti-CD28 ligated SLBs. Scale bar: 10 μm. Data are presented as Mean±SD. **G**: Western blot analysis of CD28-Y191 and Lck-Y394 phosphorylation in Lck_UD-SH3-SH2_ assisted condensation assay (see Methods). **H**-**I**: CD28 distribution (**H**) and IL-2 secretion (**I**, n=4) in Jurkat-CD28^mCherry^ cells stimulated with SEE-loaded Raji cells. Scale bar: 10 μm. Data are presented as Mean±SD.

Unexpectedly, deletion of BRS1 (CD28-dBRS1) did not completely abolish phase separation with Lck truncations, and PRS-deleted mutant (CD28-tPRS2) behaved similarly. These results suggest that terminal ICD sequences may inhibit condensate formation rather than acting passively (Figure S2C). CD28 phosphorylation (both mono- and di-phosphorylation) enhanced phase separation with Lck_UD-SH3-SH2_, wheras di-phosphorylation enhanced clustering with Lck_UD-SH3_ but abolished co-condensation with Lck_UD_ (Figure S2C-D).

To validate the role of BRS1 in cells, CD28-mBRS1^mCherry^ and CD28-mBRS2^mCherry^ were over-expressed in CD28-knockout Jurkat cells. Upon anti-CD28 stimulation, the BRS1 mutant formed fewer and smaller clusters compared with wild-type CD28 (Figure 2C and 2D). Actin polarization downstream of CD28 signaling was also impaired in CD28-mBRS1^mCherry^ cells (Figure 2E-F).

To directly assess the role of phase separation in CD28 phosphorylation, we established an in-vitro phosphorylation assay by introducing Lck-wt into pre-formed CD28-ICD/Lck_UD-SH3-SH2_ condensates and initiating phosphorylation with ATP-Mg^2+^ as previously reported (Chen et al., 2023). Lck-wt was efficiently recruited into the pre-formed condensates (Figure S3A), and immunoblotting showed reduced Y191 phosphorylation in CD28-mBRS1 compared with CD28-wt (Figure 2G). Y191 phosphorylation increased sharply 10 minutes after ATP-Mg^2+^addition, consistent with positive feedback from phosphorylation-enhanced condensation (Figure S2C-D and Figure S3B). Our previous study demonstrated that low-dose phosphorylation of CD3ε enhances its phase separation with Lck, thereby improving Lck-mediated phosphorylation efficiency (Chen et al., 2023). To examine the functional relevance of CD28/Lck phase separation in T cell activation, we over-expressed wild-type or mutated CD28 in CD28-knockout Jurkat cells and stimulated them with superantigen-loaded Raji cells (Figure 2H). The BRS1 mutation significantly impaired T cell activation, as indicated by a dramatic reduction in IL-2 secretion (Figure 2I).

### Crosstalk between TCR and CD28 during phase separation with Lck

Both CD3ε and CD28 undergo phase separation with Lck, raising the question of how their downstream signaling remains distinct. To address this question, we mixed CD3ε-ICD with either CD28-ICD or pCD28-ICD in solution and then induced phase separation by adding Lck_UD-SH3-SH2_. We observed that CD3ε-ICD and CD28-ICD were uniformly permeable within condensates, and the formed droplets exhibited dynamic behavior (Figure S4A and S4B).

To further dissect their relationship, we conducted two condensation assays: a competition mode and a recruitment mode. In the competition mode, CD3ε-ICD and CD28-ICD were pre-mixed before adding Lck_UD-SH3-SH2_. In the recruitment mode, either CD3ε-ICD or CD28-ICD were pre-mixed with Lck_UD-SH3-SH2_ to form condensates, followed by addition of the second ICD as a passenger (Figure S4C). Fluorescence intensity measurements of the passenger revealed that CD3ε-ICD /Lck condensates efficiently recruited CD28-ICD, whereas CD28-ICD/Lck condensates recruited CD3ε-ICD much less effectively (Figure S4D). Given that both CD28-ICD and CD3ε-ICD are membrane-associated in resting T in cells to prevent nonspecific activation (Dobbins et al., 2016; Xu et al., 2008; Yang et al., 2017), meaningful crosstalk likely occurs only when both receptors are co-triggered. This is consistent with previous observations that CD28 stimulation alone is insufficient to activate T cells and that TCR activation without CD28 co-stimulation yields suboptimal responses (Acuto and Michel, 2003; Lotze et al., 2024; Raychaudhuri et al., 2024).

### PD-1 specifically regulates CD28/Lck clustering

CD28 is phosphorylated by Lck by a process governed by a phase separation mechanism in which CD28/Lck clustering promotes CD28 phosphorylation, which in turn enhances condensation, forming a self-reinforcing loop. A key question is how this loop is regulated. As PD-1–SHP-2 signaling is a dominant negative regulator of CD28 (Chemnitz et al., 2004; Hui et al., 2017b), we asked whether PD-1 directly modulates CD28/Lck phase separation.

Using SLB-reconstituted CD28-ICD/Lck clusters, we found that phosphorylated PD-1-ICD (pPD-1-ICD) efficiently dissolved pre-formed pCD28-ICD/Lck_UD-SH3_ or pCD28-ICD/Lck_UD-SH3-SH2_ clusters, whereas unphosphorylated PD-1-ICD had only minimal inhibitory effect on pCD28/Lck_UD-SH3_ clusters (Figure 3A and 3B). The pPD-1-ICD mediated dissolvement was concentration dependent: at lower concentration (2 μM), pPD-1-ICD was recruited into preformed clusters but was insufficient to dissolve them (Figure S5A).

**Figure 3.**
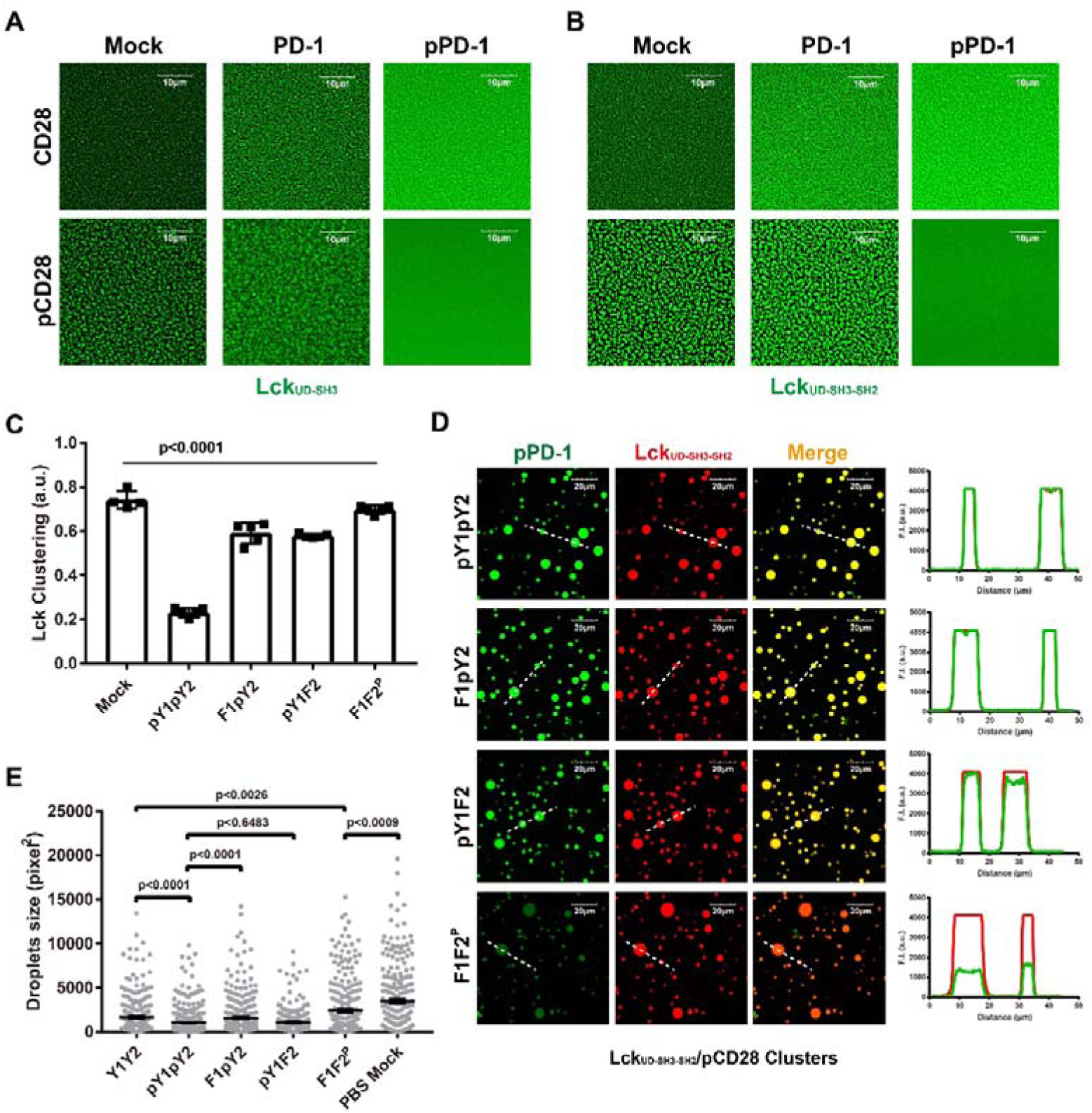
Phosphorylated PD-1 disrupts CD28/Lck condensates. **A**-**B**: Representative images showing condensates formed by Lck_UD-SH3_ (**A**) and Lck_UD-SH3-SH2_ (**B**) with wild type (upper) and phosphorylated (lower) CD28-ICD on SLB after addition of PD-1-ICD (middle) or phosphorylated PD-1-ICD (right), PBS was used as a mock control (left). Scale bar: 10 μm. **C**: Quantification of pCD28/Lck_UD-SH3-SH2_ clustering following addition of the indicated PD-1-ICDs. Data are presented as Mean±SD, n=5. **D**: Representative images of PD-1-ICD recruitment to pCD28/Lck_UD-SH3-SH2_ condensates in solution. The fluorescent intensity of AF488-labeled PD-1-ICD and AF647-labeled Lck_UD-SH3-SH2_ along the white dashed lines are shown. Scale bar: 20 μm. **E**: Droplet size distribution for pCD28-ICD/Lck_UD-SH3-SH2_ condensates in the presence of indicated PD-1-ICDs. Data are presented as Mean±SEM.

To further dissect how PD-1-ICD phosphorylation regulates pCD28-ICD/Lck phase separation, we constructed PD-1-ICD mutants carrying different tyrosine substitutions and phosphorylated them in vitro using Lck_kinase_. This produced un-phosphorylated (pPD-1-YYFF), mono-phosphorylated (pPD-1-Y1F or pPD-1-Y2F), and doubly phosphorylated (pPD-1-wt) PD-1 variants (Figure S5B). Only doubly phosphorylated PD-1 dissolved the pCD28-ICD/Lck_UD-SH3-SH2_ condensates on SLBs, whereas unphosphorylated or mono-phosphorylated PD-1-ICDs failed to do so (Figure 3C). Mono-phosphorylated PD-1-ICD, however, was recruited into pre-formed pCD28-ICD/Lck_UD-SH3-SH2_ condensates and modestly inhibited pCD28-ICD/Lck_UD-SH3-SH2_ phase separation in solution (Figure 3D-3E and Figure S5C). Notably, other co-inhibitory receptors (CTLA-4 and LAG-3) did not affect the phase separation between Lck_UD-SH3-SH2_ and CD28-ICD or pCD28-ICD (Figure S5D).

Together, these results indicate that PD-1 directly regulates CD28/Lck condensation in a phosphorylation-dependent but SHP-2-independent manner, and that this regulatory activity is highly specific among co-inhibitory receptors.

### Recruitment of PD-1 by CD28/Lck clusters requires PD-1 phosphorylation

Antigen-experienced T cells express PD-1, and we have demonstrated that PD-1 can dissolve CD28/Lck clusters independently of SHP-2. We next asked how PD-1 influences CD28 signaling during T cells re-encounter the target cells. To address this, we co-immobilized the pPD-1-ICD or PD-1-ICD with Lck_UD-SH3-SH2_ on SLBs and then introduced pCD28-ICD or CD28-ICD to mimic CD28 triggering.

Both (p)PD-1-ICD and Lck_UD-SH3-SH2_ were evenly distributed on SLBs (Figure 4A-B). The addition of CD28-ICD induced clustering of Lck_UD-SH3-SH2_ and recruited both PD-1-ICD and pPD-1-ICD (Figure S6A). In contrast, pCD28-ICD induced Lck_UD-SH3-SH2_clusters that robustly recruited pPD-1-ICD while excluding unphosphorylated PD-1-ICD (Figrue 4A-B).

**Figure 4.**
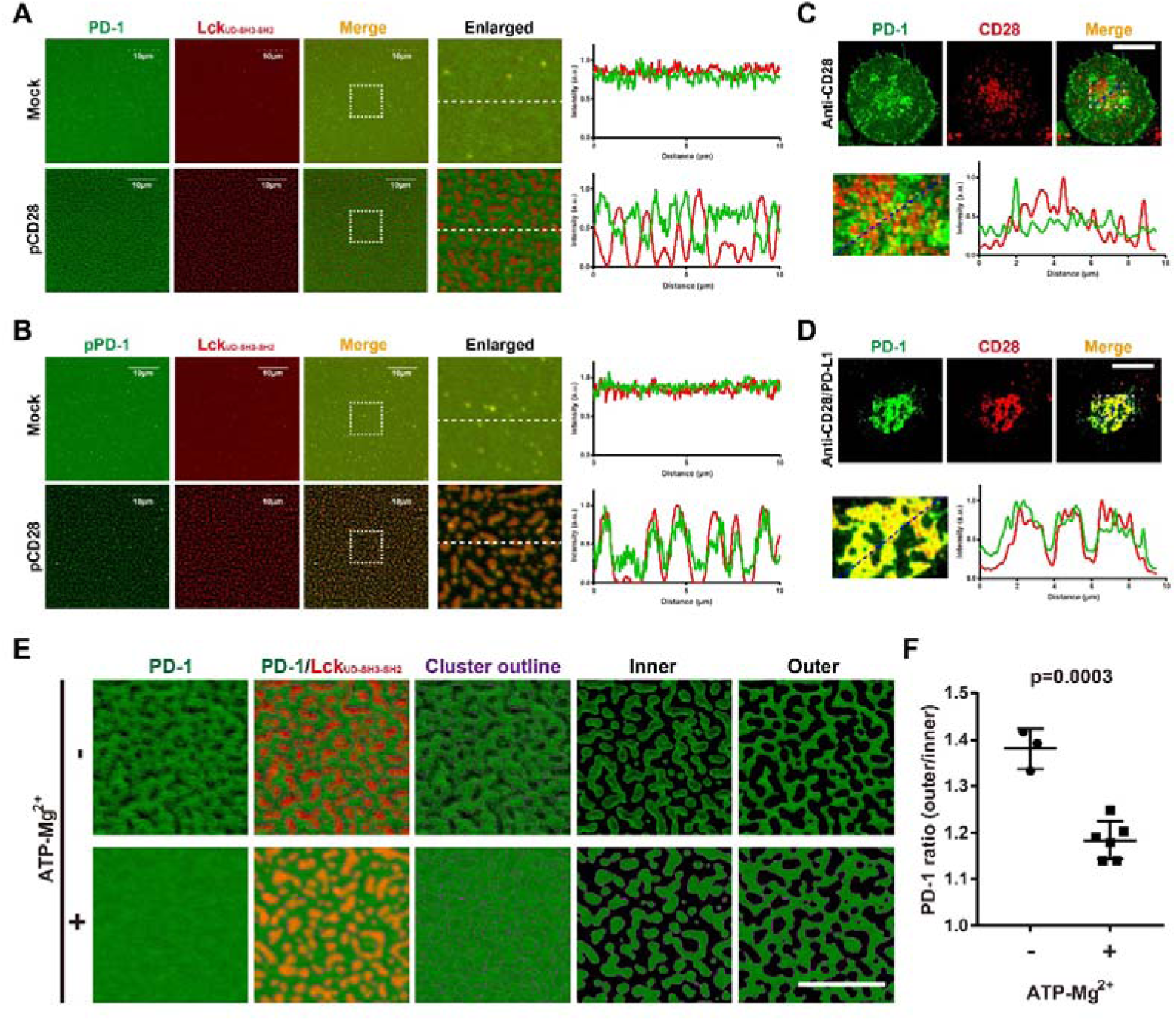
PD-1 phosphorylation controls its recruitment into CD28/Lck condensates. **A**-**B**: Localization of PD-1-ICD (**A**) and pPD-1-ICD (**B**) on SLBs upon the Lck_UD-SH3-SH2_ clustering triggered by pCD28-ICD addition. The fluorescence intensity profiles of AF488-labeled PD-1-ICD and AF647-labeled Lck_UD-SH3-SH2_ are plotted along the white dashed lines. Scale bar: 10 μm. **C**-**D**: Representative contact zones of Jurkat-CD28^mCherry^/PD-1^mGFP^ cells with anti-CD28 ligated SLBs in the presence (**C**) and absence (**D**) of PD-L1. PD-1-mGFP and CD28-mCherry intensities along the blue dashed lines are shown. Scale bar: 10 μm. **E**-**F**: Recruitment of PD-1 into pCD28-ICD/Lck_UD-SH3-SH2_/Lck-wt condensates in the absence (n=3) pr presence (n=5) of ATP-Mg^2+^. Representative images of PD-1-ICD localization on the inner and outer of the condensates are shown (**E**), the outer/inner ratio of PD-1-ICD intensity are quantified (**F**). Scale bar: 10 μm. Data are presented as Mean±SD.

To further validate these finds in cells, we overexpressed CD28 with C-terminal fused mCherry in Jurkat-PD-1^mGFP^ cells (Jurkat-CD28^mCherry^/PD-1^mGFP^) and stimulated them with immobilized anti-CD28 antibody, in the absence or presence of PD-L1 on SLBs. Anti-CD28 antibody stimulation induced CD28 clustering at the cell–SLB interface.. In the absence of PD-L1, PD-1 formed clusters at low level that were largly segregated from CD28 clusters (Figure 4C), with rear co-localized clusters, likely reflecting weak PD-1 phosphorylation induced by CD28 signaling alone. In contrast, PD-L1 engagement induced robust PD-1 phosphorylation and near-complete co-localization of PD-1 and CD28, as revealed by TIRF-SIM imaging (Figure 4D).

To rule out the effects of PD-L1 and downstream molecules such as SHP-2, we validated these results using the SLB system. We first immobilized PD-1-ICD and Lck_UD-SH3-SH2_ on SLB, followed by induction of pCD28-ICD/Lck_UD-SH3-SH2_ clustering via adding pCD28-ICD. Subsequent addition of Lck-wt, which can be recruited to these clusters, which did not initially alter PD-1-ICD exclusion. However, upon the addition of ATP-Mg^2+^, PD-1-ICD progressively penetrated into the clusters (Figure 4E and 4F).

Previous studies have shown that the phase separation of SHP-2 depends on its phosphatase domain (Zhu et al., 2020), and phosphorylated PD-1 recruits SHP-2 through its SH2 domain (Chemnitz et al., 2004; Hui et al., 2017b; Yokosuka et al., 2012). We then wondering whether SHP-2_SH2-SH2_ could phase separate with pPD-1 in the presence of Lck and thereby interfere CD28 signaling. Our results showed that SHP-2_SH2-SH2_ could not form clusters with pPD-1 on SLB, even in the presence of Lck_UD-SH3_ or Lck_UD-SH3-SH2_ (Figure S6B and S6C). Nonetheless, SHP-2_SH2-SH2_ was recruited by pPD-1-ICD, leading to the dissolvement of pCD28-ICD/Lck_UD-SH3-SH2_ clusters (Figure S6D). These results confirmed that SHP-2_SH2-SH2_ remains functional in binding PD-1 and promoting PD-1 mediated inhibition of CD28/Lck signaling, and also suggested a phosphatase-independent function of SHP2.

### PD-1 phase separates with CD28 independently of Lck

Phosphorylation of PD-1 enhances its recruitment into CD28/Lck clusters and, at high concentrations, promotes dissolution of these clusters. Because the CD28-ICD contains basic residues whereas the PD-1-IC is acidic, we asked whether PD-1 could directly interact with CD28 in the absence of Lck. If so, PD-1 may disrupt the CD28-Lck interaction through competitive binding of CD28, thereby contributing to dissolve CD28/Lck clusters.

To explore this, we immobilized phosphorylated PD-1 ICD (pPD-1-ICD) or unphosphorylated PD-1-ICD on SLBs and examined recruitment of CD28-ICD. CD28-ICD robustly phase separated with both forms of PD-1, whereas ICDs from other co-stimulatory receptors or from CD3ε did not (Figure 5A and 5B). Additionally, membrane tethering of CD28-ICD suppressed this clustering, as fewer condensates formed when His-tagged CD28-ICD was immobilized on SLB before adding His-tagged PD-1-ICD (Figure S7A).

**Figure 5.**
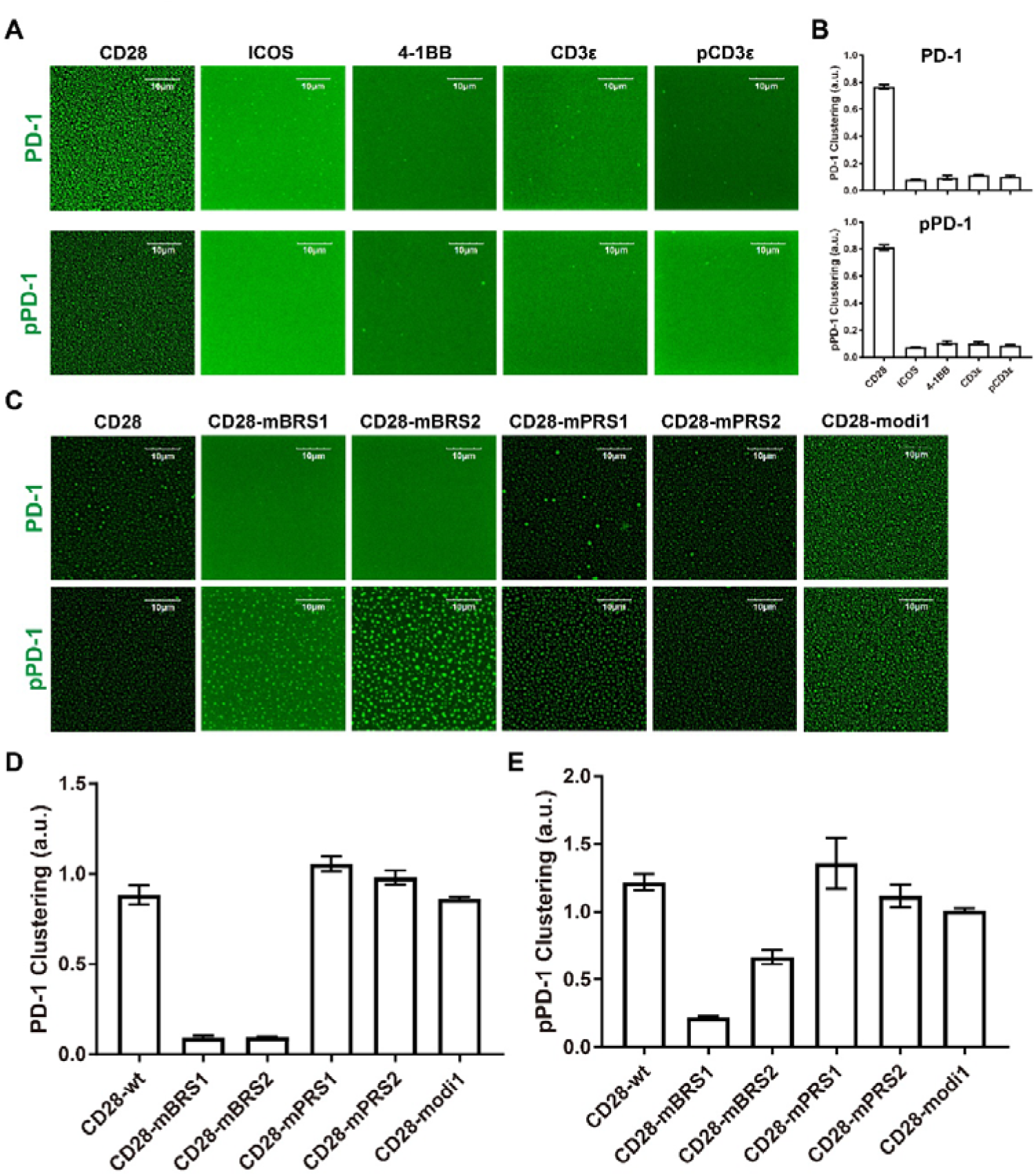
PD-1 forms phase-separated condensates with CD28. **A**-**B**: Representative images (**A**) and quantification (**B**) of PD-1-ICD (upper) and pPD-1-ICD (lower) clustering upon addition of the indicated ICDs. Scale bar: 10 μm. Data are presented as Mean±SD, n=5. **C**-**E**: Representative images (**C**) and quantification (**D** and **E**) of PD-1-ICD (**C**, upper; **D**) and pPD-1-ICD (**C**, lower; **E**) clustering with wild type or mutated CD28-ICDs. Scale bar: 10 μm. Data are presented as Mean±SD, n=5.

To further investigate the mechanism of CD28/PD-1 phase separation, we tested a series of CD28-ICD mutants. Mutation of the BRS motifs of CD28 significantly reduced its phase separation with pPD-1-ICD or PD-1-ICD (Figure 5C-5E and Figure S7B), suggesting that electrostatic interactions are a major driver of CD28/PD-1 phase separation.

It is well established that the ITSM motif of PD-1 is essential for its function and for SHP-2 recruitment (Chemnitz et al., 2004; Yokosuka et al., 2012), whereas the exact role of ITIM remains unclear. We found that PD-1 phosphorylation regulates of CD28/PD-1 phase separation, as mono-phosphorylation of PD-1 impaired its phase separation with CD28 (Figure S7C and S7D), while double-phosphorylation of PD-1 enhanced its recruitment by phase separating with CD28. These results suggest that ITIM phosphorylation may facilitate the interaction between PD-1 and CD28, contributing to CD28/PD-1 phase separation.

### CD28, PD-1, and Lck form heterogeneous clusters in cells

Phosphorylation of PD-1 not only affects its interaction with CD28 but also enables the recruitment of SHP-2, which further alters CD28/PD-1 clustering. We found that SHP-2_SH2-SH2_ could be recruited to and thus dissolve the pCD28-ICD/pPD-1-ICD clusters, but it had no effect on CD28-ICD/pPD-1-ICD clusters (Figure S8A).

Interestingly, we found that pPD-1/CD3 ε formed clusters in the presence of SHP-2_SH2-SH2_, which suggest potential interactions between pPD-1 and CD3ε (Figure S8A). The clustering dynamics of these molecules become even more complicated considering that CD28 also phase separates with Lck, which in turn efficiently phosphorylates PD-1. Using immunofluorescence, we characterized the clustering of these molecules in Jurkat-CD28^mCherry^/PD-1^mGFP^ cells stimulated with SLB presenting anti-CD28 antibody, PD-L1 or both. Lck was evenly distributed on the contact region, but its co-localization with CD28 and PD-1 was not clearly distinguishable (Figure S8B). To better resolve cluster composition, we immobilized CD28 antibody and PD-L1 on glass substrates. Under these conditions, Jurkat-CD28^mCherry^/PD-1^mGFP^ cells formed numerous clusters that were highly heterogenous in size and molecular compositions (Figure S8C). These results indicate that CD28, PD-1, and Lck dynamically regulate T cell activation by forming highly variable and adaptable clusters.

### CD28 mutations impair phase separation with PD-1 and confer resistance to ligand-independent PD-1 inhibition

TCR/CD28 signaling can induce ligand-independent PD-1 phosphorylation (Chemnitz et al., 2004), although the underlying mechanism remains unclear. Our results suggest that the triggering of TCR/CD28 activation leads to their clustering with Lck, and PD-1 can be recruited independent of ligand binding and then phosphorylated by Lck, pPD-1 in turn recruits SHP-2 to inhibit TCR/CD28 signaling. We hypothesized that CD28 mutants with reduced PD-1 binding might resist PD-1-mediated inhibition, ligand-independent as well as ligand-dependent.

To identify the key residues in CD28 for CD28/PD-1 phase separation, coarse-grained molecular dynamics (CGMD) simulations were carried out. These simulations revealed that the CD28-ICD and PD-1-ICD form dynamic condensates (Figure. 6A and Movie S1). Contact mapping identified two critical CD28-ICD regions: the RKHY motif (R203-Y206) which contacting an acidic region of PD-1 (V239-E244) via hydrophobic and electrostatic interactions (Figure. 6B-C), and a basic residue R219, which also from electrostatic interaction with the acidic residue of PD-1-ICD (Figure. 6B and 6D).

**Figure 6.**
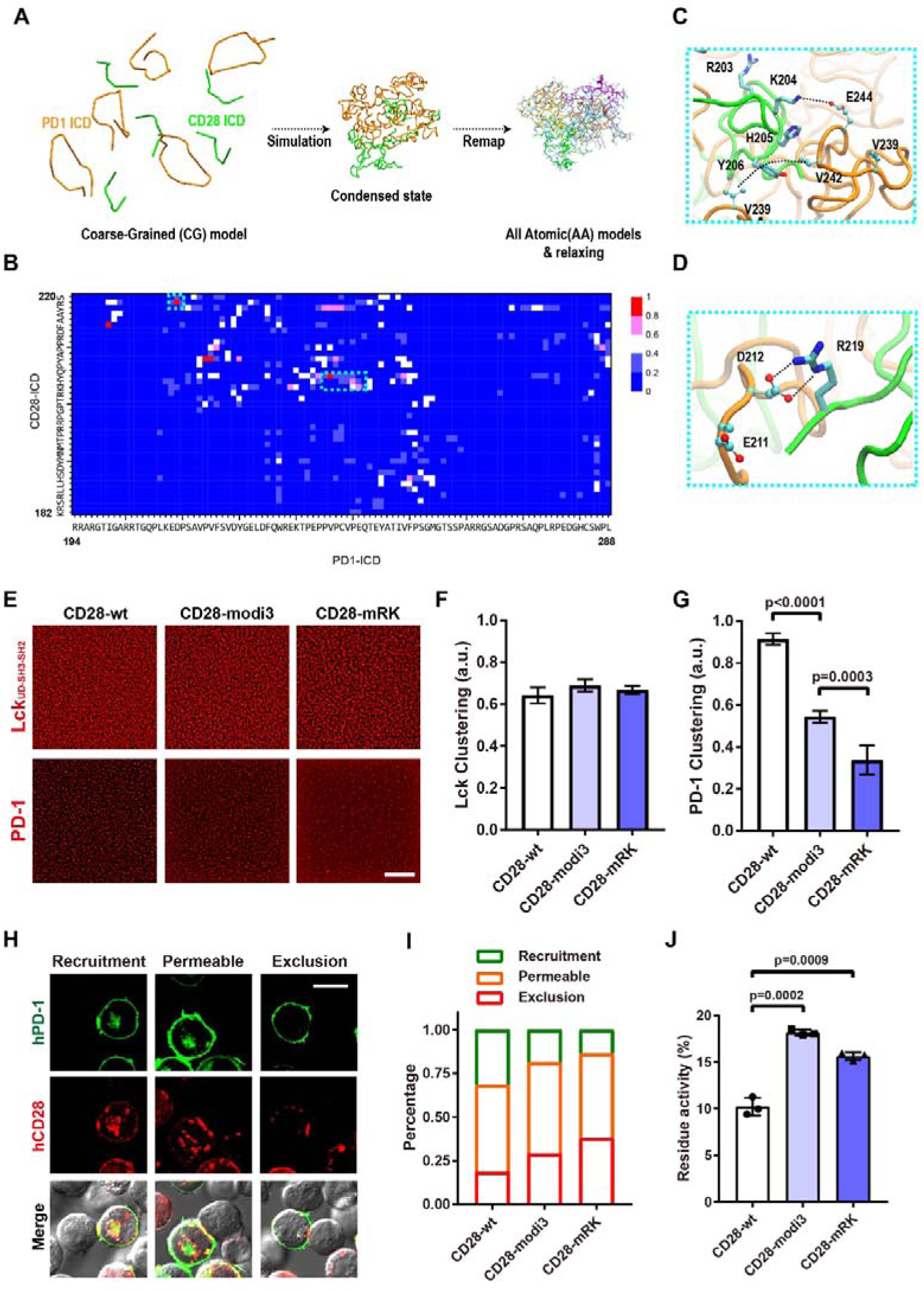
CD28 mutants disrupt PD-1 binding and confer resistance to PD-1-mediated inhibition. **A**: Workflow of CGMD simulations. Five CD28-ICDs (green) and five PD-1-ICDs (orange) adopt extended conformations (left) in a water box, condense during CGMD simulation (middle), and remapped to an all-atom model for local relaxation (right). **B**: Heatmap of CD28-ICD/PD-1-ICD residue contacts frequencies. The R219 site and the RKHY motif of CD28-ICD show a high probability interactions with PD-1-ICD (dashed lines). **C-D**: Representative all-atom structures showing interactions between the RKHY motif of CD28-ICD (green) and PD-1-ICD residues V239-E244 (orange) (**C**), and between residue R219 of CD28-ICD (green) and residue D212 of PD-1-ICD (orange) (**D**). **E**-**G**: Representative images (**E**) and quantification (**F** and **G**) of Lck_UD-SH3-SH2_ (**E**, upper; **F**) and PD-1-ICD (**E**, lower; **G**) clustering induced by wild type or mutated CD28-ICDs. Scale bar: 10 μm. Data are presented as Mean±SD, n=5. **H**-**I**: Classification of the conjugate between Jurkat-CD28^mCherry^/PD-1^mGFP^ cells and SEE-loaded Raji cells into three PD-1 localization patterns: Recruitment, Permeable and Exclusion (**H**). The distribution of conjugate types for wild type or CD28 mutant is quantified (**I**). Scale bar: 10 μm. **J**: Residual activity of Jurkat-CD28^mCherry^/PD-1^mGFP^ cells stimulated with 20 ng/ml SEE-loaded Raji cells upon PD-1 triggering. The residue activity is calculated as (IL2-IL2^PD-L1^)/IL2, where IL2 and IL2^PD-L1^ refers to the raw IL-2 secretion of Jurkat-CD28^mCherry^/PD-1^mGFP^ cells stimulated with PD-L1 negative or positive Raji cells, respectively. Data are presented as Mean±SD, n=3.

CD28 mutants designed based on these predictions (CD28-modi3 and CD28-mRK) showed impaired condensation with PD-1 while retaining normal phase separation with Lck_UD-SH3-SH2_ (Figure 6E-6G). Upon phosphorylation, pCD28-mRK lost the ability to phase separate with Lck, whereas both pCD28-modi3 and pCD28-mRK showed reduced clustering with PD-1 (Figure S9A-D) and resisted PD-1–mediated dissolution of CD28/Lck condensates (Figure S9E-F).

We next investigated the recruitment of PD-1 by CD28 mutants in activated Jurkat cells, Jurkat-CD28^mCherry^/PD-1^mGFP^ cells were stimulated with SEE loaded Raji cells which did not express PD-L1 or PD-L2. CD28 formed clusters at the center of Jurkat-Raji contact interface, whereas PD-1 localization could be classified into three patterns: Recruitment, in which PD-1 was enriched within CD28 clusters; Permeable, in which PD-1 was distributed evenly; and Exclusion, in which PD-1 was excluded and separated from CD28 clusters (Figure 6H). CD28-modi3 and CD28-mRK significantly increased PD-1 exclusion and reduced PD-1 recruitment as shown by increased ratio of Exclusion and decreased ratio of Recruitment compared with CD28-wt (Figure 6I). Functionally, Jurkat cells expressing CD28-modi3 and CD28-mRK produced less IL-2 in the absence of PD-L1 compared to cells expressing CD28-wt (Figure S9G). In contrast, when stimulated with PD-L1 expressing Raji cells, Jurkat cells expressing CD28-modi3 or CD28-mRK secreted more IL-2 than CD28-wt cells, although this difference diminished at high SEE concentrations (Figure 6J and Figure S9G-S9H).

### Engineering CD28 mutations into 2^nd^ CAR-T cells enhances resistance to PD-1 inhibition

We next compared the phase separation preferences of CD28 variants with Lck_UD-SH3-SH2_, PD-1 or pPD-1. CD28 mutations exhibited distinct condensation preferences. Phosphorylation of CD28-ICD enhanced phase separation with Lck_UD-SH3-SH2_ but suppressed condensation with PD-1-ICD and pPD-1-ICD. A similar trend was observed for BRS mutations, whereas PRS mutations showed the opposite behavior (Figure 7A).

**Figure 7.**
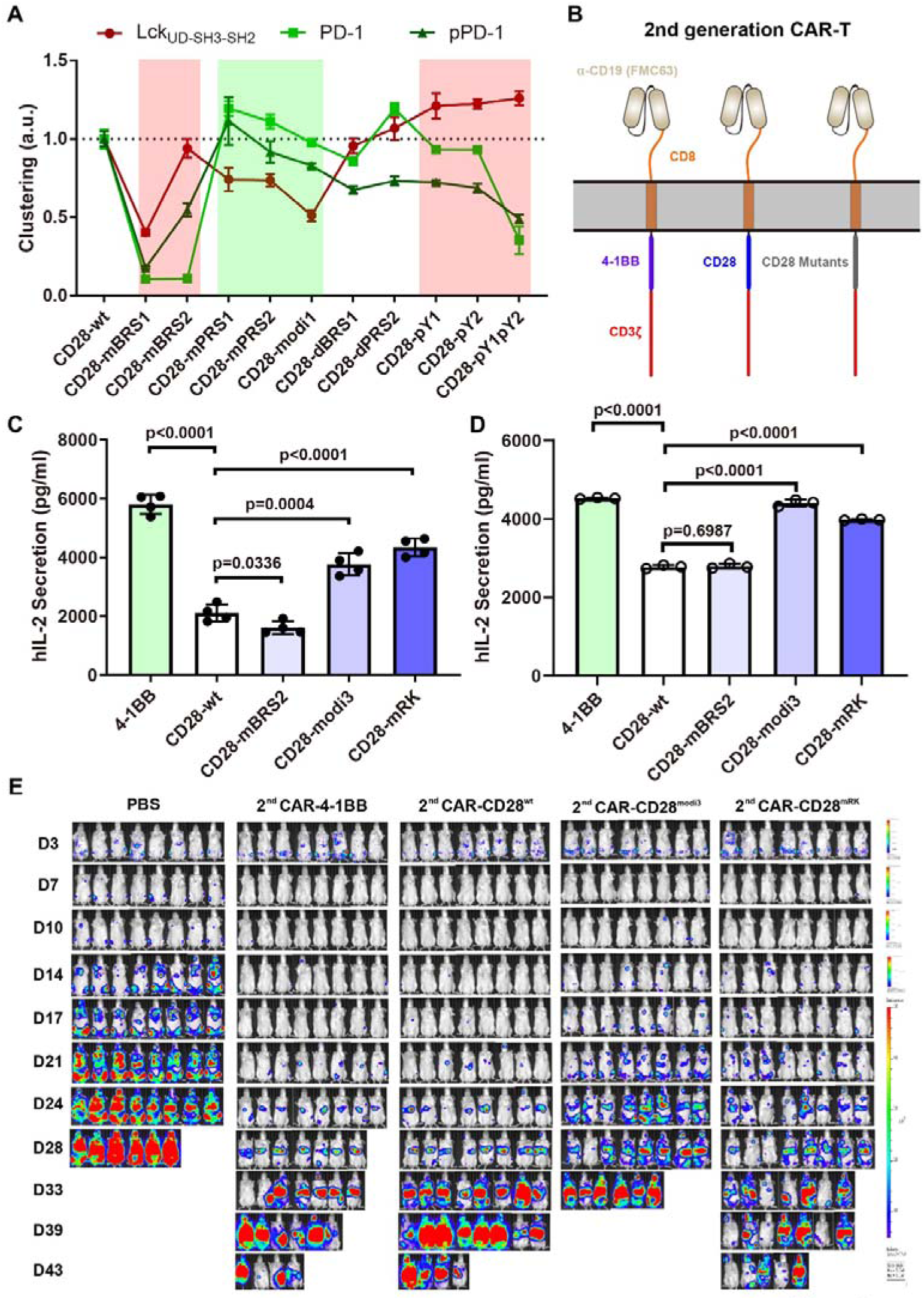
CD28 mutants incorporated into CAR-T cells enhance antitumor activity. **A**: Quantification of Lck_UD-SH3-SH3_, PD-1-ICD and pPD-1-ICD clustering triggered by wild type, mutant or phosphorylated CD28-ICDs. Data are presented as Mean±SD, n=5. **B**: Schematic of the second-generation CAR-T design used in this study. **C**-**D**: IL-2 secretion by second-generation CAR-Jurkat cells stimulated with PD-L1 expressed Raji cells in the absence (**C**, n=4) or presence (**D**, n=3) of anti-PD-L1 antibody. Data are presented as Mean±SD. **E**: Bioluminescence imaging of Nalm6-ffluc tumor burden in NSG mice treated with indicated second-generation CAR-T variant cells.

CD28 variants that preferentially phase separated with Lck while exhibiting reduced condensation with (p)PD-1 were selected for CAR engineering (Figure 7B). Several second-generation 28z CARs were transduced into Jurkat cells. Most constructs displayed comparable surface expression, except for CD28-mBRS1, which failed to localize to the membrane (Figure S10A and S10B). None of the CAR constructs exhibited tonic signaling, as indicated by minimal IL-2 secretion in the absence of target cells (Figure S10C).

Upon stimulation with Raji cells, all engineered Jurkat cells were robustly activated, although BRS2 mutations slightly reduced IL-2 secretion (Figure S10D). When stimulated with PD-L1–expressing Raji cells, Jurkat cells expressing wild-type CD28-CAR or BRS2-CAR showed impaired IL-2 production, whereas cells expressing 4-1BB–CAR, CD28-modi3-CAR, or CD28-mRK-CAR retained functional activity (Figure 7C and S10E). Blockade of PD-1 signaling with anti–PD-L1 antibody revealed that CD28-modi3 and CD28-mRK CARs were less sensitive to ligand-independent PD-1 inhibition (Figure 7D).

Finally, in a murine xenograft B cell lymphoma mode, CD28-mRK significantly improved the antitumor efficacy, achieving tumor control and survival comparable to that of 4-1BB-based CAR-T cells (Figure 7E and Figure S10F).

## Discussion

Co-stimulatory receptors provide essential secondary signals that enable productive T cell activation and have been widely adopted in engineered T cell therapies. Despite decades of biochemical and functional characterization, how CD28 signaling is physically organized and dynamically regulated has remained poorly defined. In this study, we identify phase separation as a central mechanism governing CD28 co-stimulation and its suppression by PD-1, thereby extending condensate-based signaling principles beyond the TCR to the co-signaling network. Incorporation of this competitive phase-separation mechanism into CAR-T construct enhances antitumor efficacy.

### CD28 as a phase-separating co-signaling receptor within a broader signaling landscape

In resting T cells, the CD28 ICD is sequestered against the inner leaflet of the plasma membrane through electrostatic interactions mediated by basic-rich sequences, maintaining CD28 in a signaling-repressed state (Dobbins et al., 2016; Yang et al., 2017). Our data reveal that ligand engagement releases the CD28 ICD, enabling phase separation with Lck and formation of signaling-competent condensates. These condensates amplify CD28 phosphorylation and downstream signaling through a positive feedback loop, analogous to previously described TCR/Lck and LAT-based assemblies. Notably, while the PRS of CD28 is known to interact with Lck SH3 domains, our results indicate that membrane release via basic-rich motifs, rather than SH3-mediated binding alone is the dominant determinant of CD28/Lck condensation. Similar observations have been made in the CD3ε/Lck condensation system (Chen et al., 2023; Holdorf et al., 1999).

We further found that other co-signaling receptors, such as 4-1BB and ICOS, do not undergo phase separate with Lck under comparable conditions. However, these receptors may instead engage other Src -family kinases. For example, Fyn kinase has been reported to phosphorylate CD3ε, CD3ζ and CD28 in vitro and in vivo (Holdorf et al., 1999; Li et al., 2017; Raab et al., 1995). Whether distinct Src-family kinase form receptor-specific condensates remains an open question and will require further systematic investigation.

### PD-1 disrupts CD28/Lck condensation through multi-layered mechanisms

PD-1 is well established as a potent inhibitor of CD28 signaling through recruitment of SHP-2 (Hui et al., 2017a; Kamphorst et al., 2017). Unexpectedly, we find that phosphorylated PD-1 can directly dissolve CD28/Lck condensates independently of SHP-2, revealing an additional layer of regulation. Mono-phosphorylated PD-1 partitions into CD28/Lck condensates, whereas dual phosphorylation of PD-1 actively destabilizes these assemblies through competitive binding to the CD28 ICD. This phosphorylation-dependent, multistate behavior provides a mechanistic explanation for graded PD-1 inhibition and helps reconcile prior observations that PD-1 function is only partially dependent on phosphatase recruitment.

Moreover, CD28 and PD-1 ICDs can phase separate directly, even in the absence of Lck, suggesting that once CD28 is released from the membrane it occupies a regulatory bifurcation point. At this juncture, CD28 may nucleate activating CD28/Lck condensates or inhibitory CD28/PD-1 assemblies, with the balance between these states dynamically tuned by kinase and phosphatase activities. This competitive condensate model offers a physical framework for understanding ligand-independent PD-1 activity and the incomplete restoration of T cell function observed following checkpoint blockade.

### Condensate selectivity as an engineering principle

The dual ability of CD28 to phase separate with both Lck and PD-1 reveals a new dimension of co-signaling regulation, that is, competition for condensate incorporation. Decoupling CD28 from PD-1 while maintaining its ability to engage Lck provides a rational strategy for enhancing engineered T cells. We identified CD28 mutants (CD28-modi3 and CD28-mRK) that preferentially form with Lck but not PD-1. These mutants enhanced cytokine production in Jurkat cells, strengthened signaling in second-generation CAR constructs, and improved CAR-T cell antitumor efficacy in vivo.

This framework provides a biophysically grounded rationale for why CD28-derived co-stimulatory domains have been successfully used across CAR-T platforms and why modifications that modulate condensate behavior can dramatically tune therapeutic performance.

### Implications for the broader family of co-signaling receptors

Recent efforts to incorporate co-signaling domains into engineered T cells, including TCR-T, TRuC-T, STAR-T/HIT-T and Co-STAR systems, underscore the fundamental importance of CD28 signaling in therapeutic T cell function (Dobrin et al., 2024; McCarthy et al., 2023; Mog et al., 2024). The mechanistic insights provide here suggest that successful co-signaling engineering may depend on selecting domains that not only activate canonical signaling cascades but also form appropriate condensates in response to antigen engagement, such as use BiTS to drive Lag-3 and TCR into close proximity for autoimmune disease (Du et al., 2025).

As more co-stimulatory and co-inhibitory receptors are examined through the lens of phase separation, it is likely that phase separation will emerge as a common mechanistic theme underling their activation thresholds, signal tuning capacities and susceptibility to checkpoint inhibition.

## Concluding remarks

In summary, our study reveals phase separation as a fundamental mechanism regulating CD28 co-stimulation and its suppression by PD-1. By uncovering a competitive condensate-based regulatory logic and leveraging it for rational engineering, we establish condensate selectivity as a new principle for the design of next-generation T cell therapies.

## Data Availability Statement

All materials generated or analyzed in this study are included in this published article and its Supplementary Information document. For special materials, please contact the lead contact. Any material that can be shared will be released for noncommercial use through a material transfer agreement.

## Acknowledgement

We would like to thank Prof. Chenqi Xu for for useful comments and discussions. We thank Dr. Yun Feng and Qing Bian for the help in TIRF-SIM imaging and data processing with Imaris. We thank J. Jia and S. Meng (Core Facility, Institute of Biophysics, CAS) for technical support in the flow cytometry analysis. All the imaging experiments were performed at the Center for Biological Imaging (CBI), Institute of Biophysics, Chinese Academy of Science. The computational resources in this study were provided by the Harbin Supercomputer Center and HPC-Service Station at the CBI of the Institute of Biophysics.

## Author contribution

H. C. and J. L. initiated the project, and H. C., C. L., W. C. and J. L. conceived the project and designed the experiments. H. C. constructed the cell lines, performed the phase separation assay and cell function experiments. S. W. performed mouse tumor studies. Y. Z. performed MD simulations and analyzed the results. H. C. and W. H. purified the proteins. H. C. and J. L. prepared initial figures and manuscript. W.C., and C.L. participated in the discussions and optimized figures. All authors helped revise the manuscript.

