## Supporting materials for "CD28 Co-stimulation Is Organized by Signaling Condensates and Counteracted by PD-1"

**Materials and Methods**

**Antibodies**

Antibodies used in this study were purchased from various commercial suppliers: Biotinylated anti-CD3ε (UCHT1, Abcam, ab191112); PE labelled anti-Lck (pY505, BD, 558552); Anti-phospho-tyrosine (4G10, Millipore, 05-321); Goat anti-Rabbit IgG-APC (Solarbio, K0034G-APC); anti-CD3 (eBioscience, 16-0037-85); APC labelled anti-CD3 (eBioscience, 17-0038-42); biotinylated anti-CD28 (Abcam, ab267385); anti-CD28 (eBioscience, 16-0289); anti-phospho-CD28 (Tyr191) (Cell Signling Technology, 16399S); anti-PD-L1 (ABclonal, A20270); SEE (SAB, AP70996).

**Cell lines and lentiviral transfection**

Lentiviral constructs were co-transfected into HEK293T cells with helper plasmids pMD2.G and psPAX2 using Polyplus transfection reagent (Polyplus, #101000046). Viral supernatant was harvested at 48 and 72 h of transfection.

For CD28 clustering assays, wild-type or mutant CD28-mCherry constructs were reconstituted into CD28 KO Jurkat cells or Jurkat^PD1-mGFP^ cells ( gift from Dr. Guangshuo Ou, Tsinghua University), followed by flow cytometry sorting. These Jurkat derivatives were also used for CD28 localization analysis.

CAR-Jurkat cells was generated by infecting Jurkat cells with lentivirus encoding CAR constructs, followed by quantification of surface CAR expression using anti-Mouse Fab-Alexa647 (Jackson Immunoresearch, #115-606-072) and flow cytometry.

All Jurkat cells were cultured in RPMI-1640 medium supplemented with 100 U/mL streptomycin, 100 ug/mL penicillin and 10% fetal bovine serum at 37℃ with 5% CO_2_.

**Peptides and proteins**

*Synthesized peptides*

All peptides used in this study were synthesized by GL biotech (Shanghai). Fluorescence dye labeling was conducted as previously described (Chen et al., 2023).

*Intracellular domain of PD-1*

The human PD-1 intracellular domain (residues 192-288) was cloned into a home modified pCOLD vector encoding an N-terminal GST tag followed by an 8×His-tag and a 3C protease cleavage site. PD-1-ICD mutations were introduced by overlapping PCR. Constructs were expressed in BL21 (DE3) *E. Coli* (Tsingke).

Cells were harvested and lysed, supernatant were collected and then incubated with GST-tag purification resin. GST-tag were removed by 3C protease digestion. The effluents were collected and boiled, supernatants were further purified by gel filtration (Hiload 16/600, Superdex 200, Cytia).

For phosphorylation, purified PD-1-ICD was incubated with Lck_kinase_ domain in the presence of ATP-Mg^2+^, followed by removal of Lck_kinase_ and excess ATP by gel filtration (Hiload 16/600, Superdex 200, Cytia). The phosphorylation was confirmed by mass spectrometry.

*Lck proteins*

The expression and purification of Lck proteins were conducted as previously described (Chen et al., 2023). Briefly, full-length Lck constructs (Lck-wt: residues 3-509, and Lck-open: residues 3-509 with K273R/Y505F mutation) were expressed in HEK293T cells. The kinase domain of Lck (Lck_kinase_: residues 245-501) was produced in BTI-Tn-5B1-4 insect cells. Lck regulatory fragments (Lck_UD_: residues 3-60; Lck_UD-SH3_: residues 3-121; Lck_UD-SH3-SH2_: residues 3-226) were expressed in BL21 (DE3) *E. Coli* (Tsingke). All Lck proteins were purified with His-Trap HP column followed by gel filtration (Hiload 16/600, Superdex 200, Cytia).

*SHP-2 protein*

The SH2 domain of SHP-2 (residues 1-247) was cloned with an N-terminal GST-tag followed by 3C cleavage site and C-terminal 8×His-tag. Protein expression was induced in BL21 (DE3) *E. Coli* (Tsingke) with IPTG. Cells were harvested and then lysed in PBS buffer with 2 mM DTT and 1× protease inhibitor cocktail supplement. Supernatant was subjected into GST-tag purification resin; GST-tag was removed by 3C protease digestion. The effluent was collected and purified by gel filtration (Hiload 16/600, Superdex 200, Cytia).

*Extracellular domain of PD-L1*

The PD-L1 extracellular domain (ECD) was fused with a C-terminal AVI-tag and followed by an 8×His-tag, subcloned into a home-modified phage-vector. The recombined plasmid was transduced into Expi-293F cells for protein expression. Supernatants were collected and purified by His-Trap HP column, proteins were then biotinylated with BirA and further purified by gel filtration (Hiload 16/600, Superdex 200, Cytia).

**Protein condensation assays on supported lipid bilayer (SLB)**

SLB formation and protein condensation assays were performed as previously described (Du et al., 2025). Briefly, small unilamellar vesicles (SUV) were applied to pre-cleaned glass-bottom 96-well plates and incubated at 37 ℃ for 1 hour to form bilayers. His-tagged proteins were ligated to the pre-formed SLBs, and the passenger molecules were added to trigger on-membrane phase separation. Imaging were conducted using an Olympus FV1200 microscope equipped with a 60× oil immersion objective.

**CAR-T cell preparation**

Lentiviral particles contained CAR constructs were harvested and concentrated, and used to infect the target cells in the presence of 8 µg mL^−1^ Polybrene, followed by spinoculation at 1600 rpm for 60 min. CAR surface expression was validated using anti-Mouse Fab-Alexa647 (Jackson Immunoresearch, #115-606-072) and flow cytometry.

For primary CAR-T cells, pan CD3^+^ T cells were isolated from PBMCs (Milestone® Biotechnologies) using the MojoSort Human CD3 T cell isolation Kit (Biolegend, #480022). T cells were activated with Human T-activator CD3/CD28 Dynabeads (Thermo Fisher Scientific, #11161D) on day 0, transduced with CAR lentivirus at day 1, and expanded with media doubling and human IL-2 supplementation every 2-3 days. Dynabeads were removed on day 7 with magnetic separation. CAR-T cells were harvested and the transduction efficiency was assessed by ZsGreen and CAR surface staining.

**Jurkat-Raji co-culture**

For IL-2 secretion assays, 2×10^5^ SEE-loaded Raji cells or Raji^PDL1-mCherry^ cells were co-cultured with 1×10^5^ Jurkat^CD28-mCherry^ cells in DMEM medium for 18-24 hours. Supernatants was collected and analyzed using ELISA kit for IL-2 quantification.

For CAR-Jurkat cells, 2×10^5^ Raji cells or Raji^PDL1-mCherry^ cells were co-cultured with 1×10^5^ CAR-Jurkat cells in the absence or presence of anti PD-L1 antibody.

For imaging of Jurkat-Raji conjugates, SEE loaded Raji cells and Jurkat cells were plated in optical bottom 96-well plates and imaged with Olympus FV1200 confocal microscope equipped with a 60× oil immersion objective.

**TIRF-SIM imaging**

Co-localization of CD28/Lck, PD-1/CD28 and CD28 clustering were quantified with TIRF-SIM imaging as previously reported (Chen et al., 2023). In brief, divalent streptavidin was attached to the biotin-ligated SLB or glass surface, followed by biotinylated CD28 antibody or PD-L1 ligation. Jurkat cells which stably expressing wild-type or mutated CD28 and PD-1 were added and incubated at 37℃ for 40 min. Cells were then fixed and imaged with a TIRF-SIM microscope. Images processing and data analyses were carried out with Image J and Imaris 10 (Bitplane, AG).

**Molecular dynamic simulation**

All-atom models of human CD28-ICD (residues K182 to S220) and human PD-1-ICD (residues R194-L288) were converted to coarse-grained (CG) MARTINI models using martinize.py (http://cgmartini.nl). Five copies of each were placed randomly and well-separated in a ~30×30×30 nm^3^ rectangular water box with 0.15 M KCl. Energy minimization was employed using both the steepest-descent and conjugate gradient algorithms (maximum force < 100 kJ/mol nm). Subsequently, a 12 µs CGMD simulation without any elastic restrains was conducted

.

Simulations used a 310K velocity-rescale thermostat with a coupling constant of 1 ps and 1 atm Parrinello-Rahman barostat with a coupling constant of 6 ps. Verlet cutoffs scheme with automated buffered tolerance was employed to generate the pair list for calculating non-bonded forces. Long-range non-bonded interactions were calculated using the Reaction Field method. A potential shift was applied to the Coulomb and Lennard-Jones (LJ) interactions which shifts the potential by a constant such that the potential is zero at a cutoff of 1.2 nm. The time step was set to 25 fs and the snapshots were recorded every 5 ns. All CGMD simulations were performed using the MARTINI coarse-grained force field under periodic boundary conditions as implemented in the GROMACS-2023 software package. Contact maps was computed from the last 9 µs trajectory using a 0.6 nm residue-residue distance cutoff. between the sidechain beads of most residues (backbone bead for Gly and Ala residue).

To observe the inter-residue interactions at the atomic level, an 8µs CG snapshot was back-mapped to an all-atom model. This model was solvated in rectangular water box using the TIP3P water model with a charge neutrality and ~150 mM salt concentration. Equilibration was carried out, followed by production simulations of ~100 ns. Temperature was controlled at 310 K with Langevin dynamics and pressure was maintained at 1 atm with the Nosé-Hoover Langevin piston method. Particle Ewald Mesh summation was used for electrostatic calculation and a 12 Å cutoff was used for short-range non-bounded interactions. Energy minimizations and molecular dynamics simulations were performed with NAMD using the CHARM36m force field for proteins under periodic boundary conditions.

**Mouse Experiments**

Female NSG mice (6-8weeks, Beijing HFK Bioscience Corporation) were housed in the animal facility of the Institute of Biophysics, Chinese Academy of Sciences. Mice were injected intravenously with 0.5million Nalm6-firefly luciferase-p2A-mCherry cells on day 0 and treated with 1 million CAR-T cells intravenously on day 4. Disease progression was monitored by bioluminescence imaging and survival analysis.

For in vivo phenotyping studies, 2.5 µg human recombinant IL-2 was administered subcutaneously every 2 days. All procedures were approved by the Biomedical Research Ethics Committee of the Institute of Biophysics, Chinese Academy of Sciences (SYXK2023189), and conducted according to the institutional guidelines.

Chen, H., Xu, X., Hu, W., Wu, S., Xiao, J., Wu, P., Wang, X., Han, X., Zhang, Y., Zhang, Y.*, et al.* (2023). Self-programmed dynamics of T cell receptor condensation. Proc Natl Acad Sci U S A *120*, e2217301120.

Du, J.S., Chen, H., You, J., Hu, W., Liu, J., Lu, Q., Zhang, Y., Gao, J., Lin, M.J., Foster, C.J.R.*, et al.* (2025). Proximity between LAG-3 and the T cell receptor guides suppression of T cell activation and autoimmunity. Cell *188*, 4025-4042.

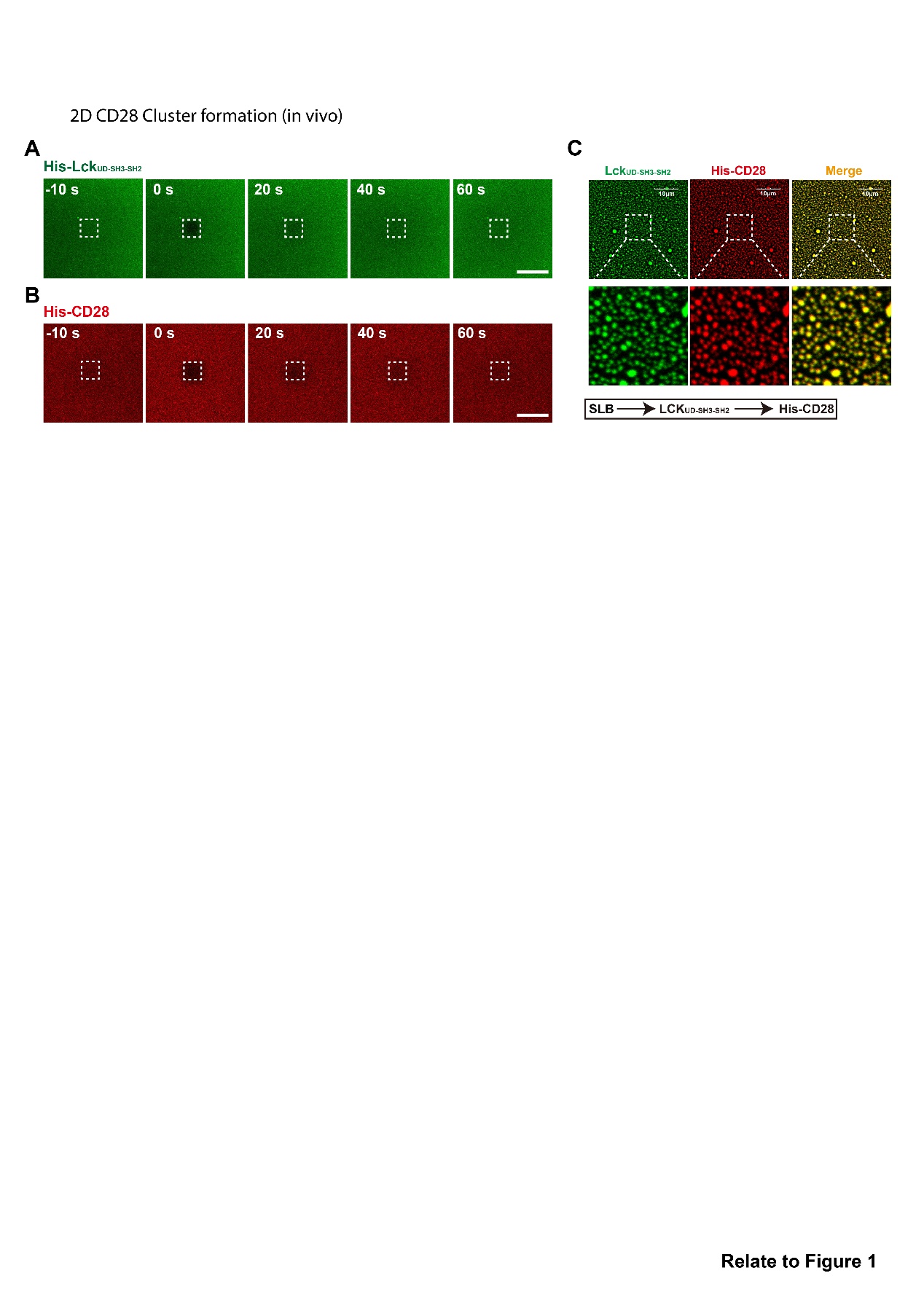

**Figure S1. CD28 and Lck form co-localized phase separated condensates on SLBs.**

**A-B**: Time-lapse snapshots of AF488-labeled Lck_UD-SH3-SH2_ (**A**) and AF647-labeled CD28-ICD (**B**) fluorescence recovery after photobleaching on SLBs. Scale bar: 10 μm.

**C**: Representative image of CD28-ICD/Lck_UD-SH3-SH2_ phase separation on SLBs with the indicated protein-addition sequence. Scale bar: 10 μm.

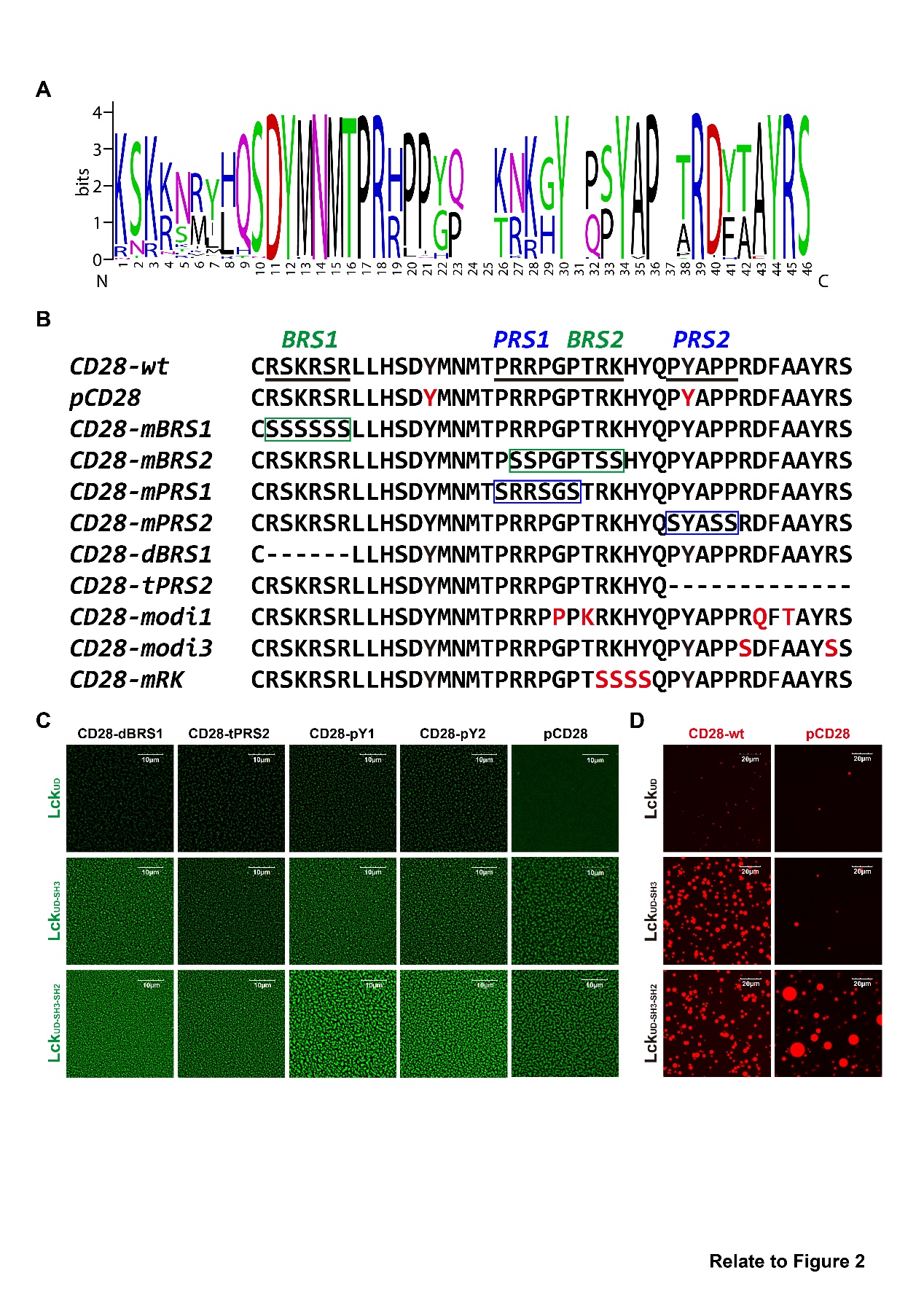

**Figure S2. Sequence regions governing CD28/Lck condensation.**

**A:** Sequence conservation analysis of CD28-ICD.

**B:** CD28-ICD constructs used for phase separation assays, phosphorylated tyrosines are marked in red.

**C-D:** Representative images showing condensates formed by CD28-ICD mutants and Lck truncates on SLBs (**C**) or in solution (**D**). Scale bars: 10 μm (**C**) and 20 μm (**D**).

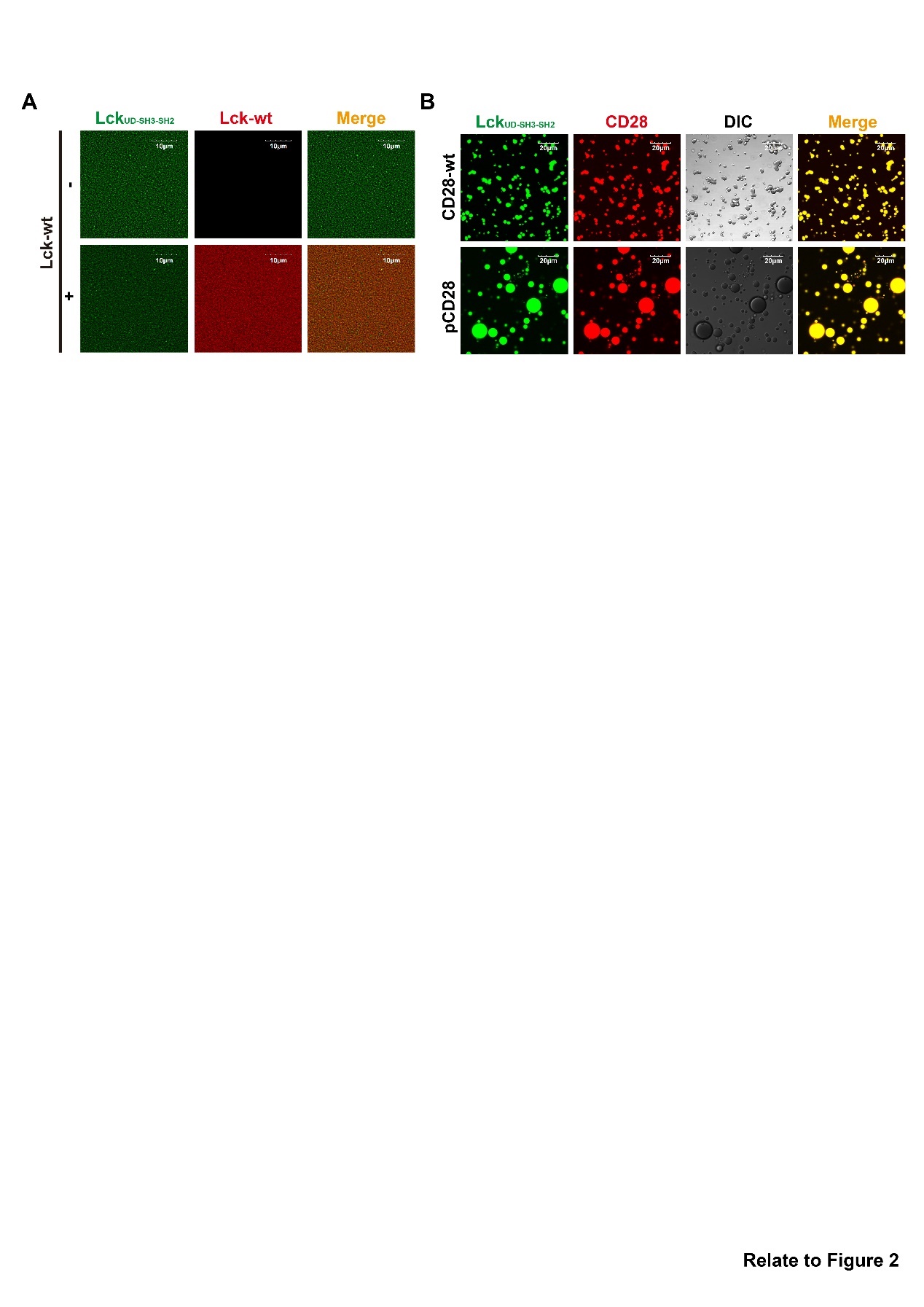

**Figure S3. CD28/Lck condensates recruit Lck and are enhanced by CD28 phosphorylation.**

**A**: Recruitment of AF647-labeled Lck-wt into CD28-ICD/Lck_UD-SH3-SH2_ (AF488 labeled) condensates. Scale bar: 10 μm.

**B**: Condensates formed by Lck_UD-SH3-SH2_ (AF488 labeled) with wild type and phosphorylated AF647-labeled CD28-ICD in solution. Scale bar: 20 μm.

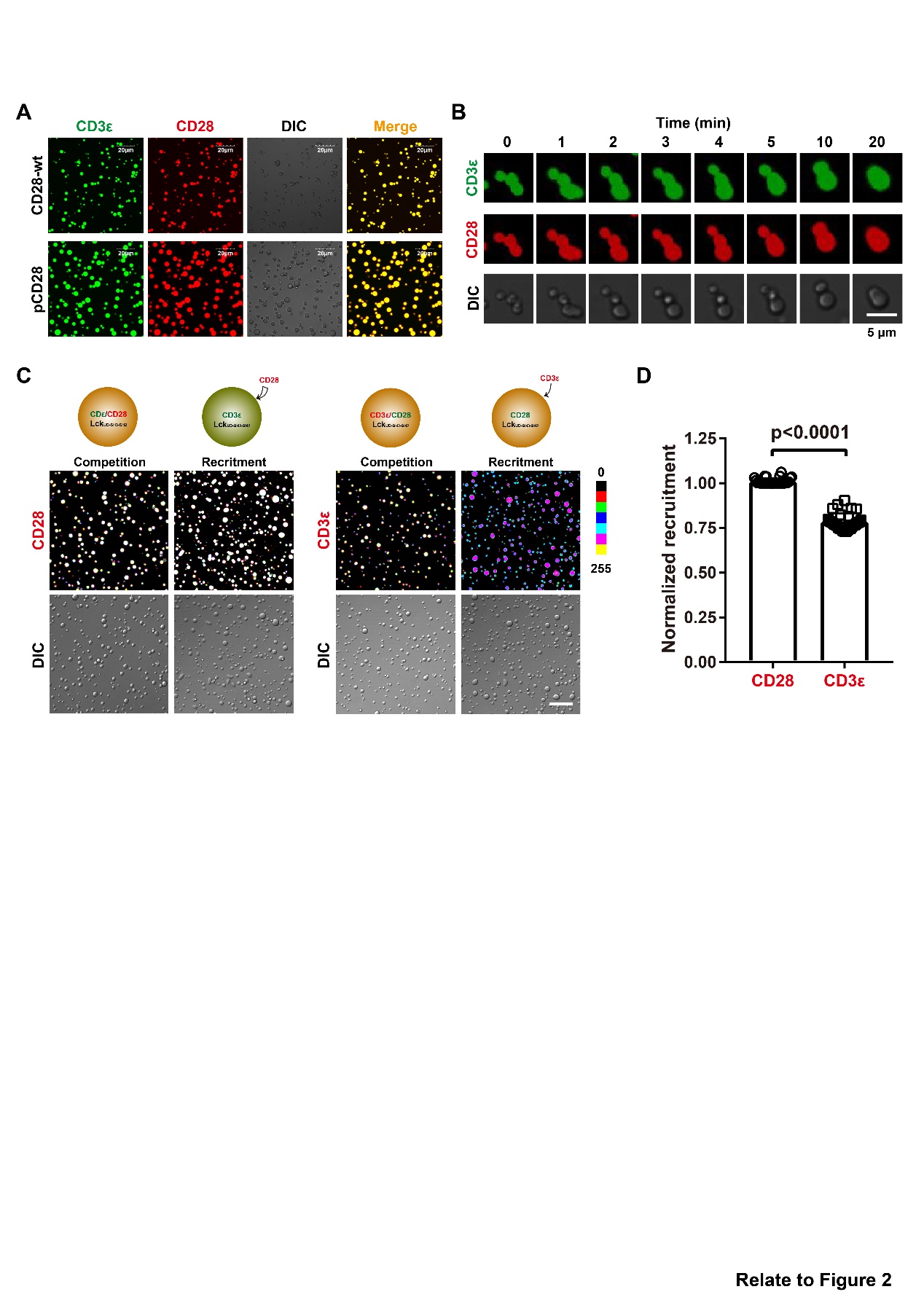

**Figure S4. Crosstalk between CD28/Lck and CD3ε/Lck condensates.**

**A**: Condensates formed by Lck_UD-SH3-SH2_ with wild type and phosphorylated CD28-ICD (AF647 labeled) in the presence of CD3ε-ICD (AF488 labeled). Scale bar: 20 μm.

**B**: Time-lapse snapshots of CD3ε/CD28/Lck_UD-SH3-SH2_ condensates fusion in solution. Scale bar: 5 μm.

**C-D**: Representative images (**C**, lower) and quantification (**D**, n=30) of CD28-ICD and CD3ε-ICD recruitment, the schematic of competition and recruitment assays were shown (**C**, upper). Scale bar: 20 μm. Data are presented as Mean±SD.

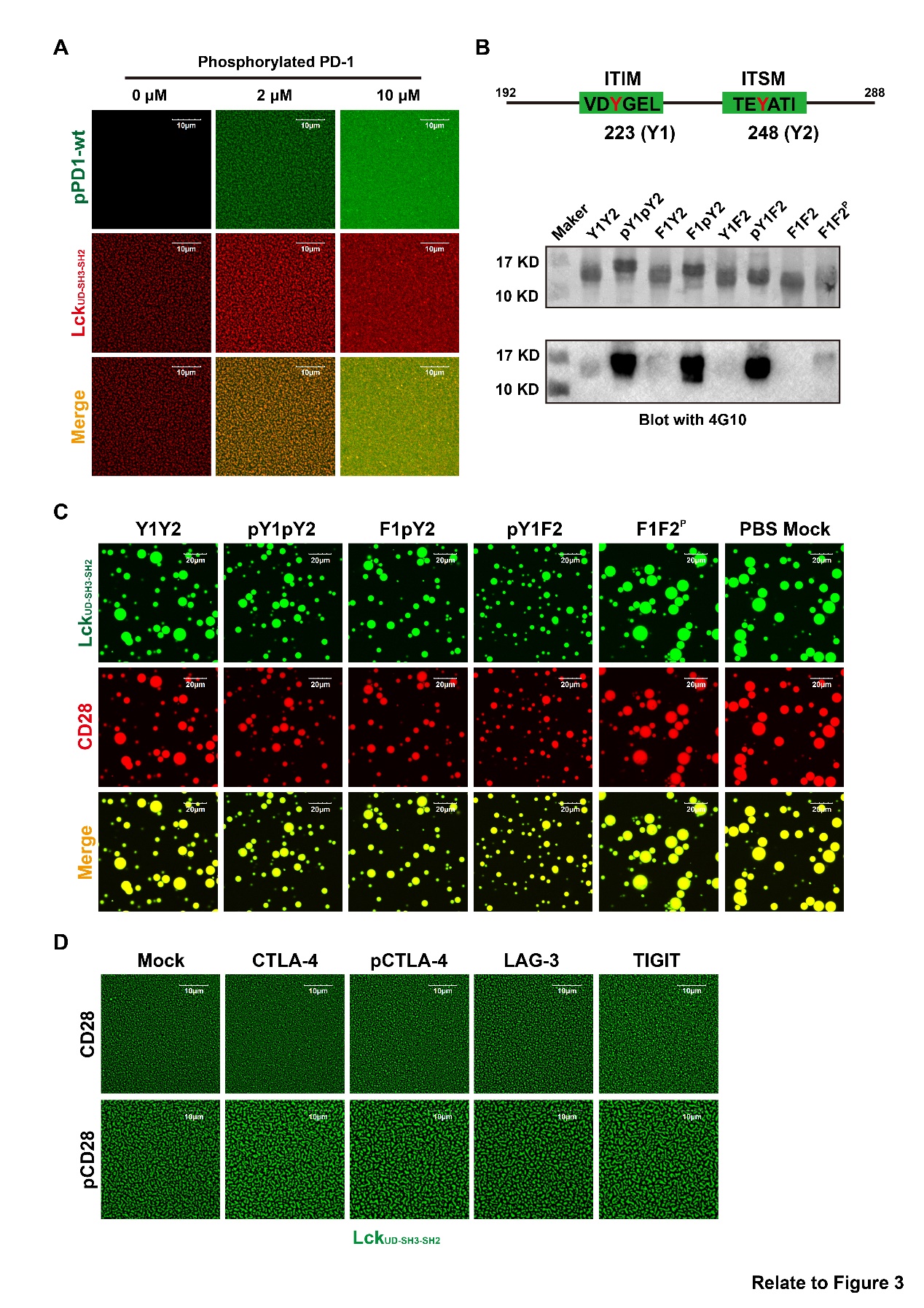

**Figure S5. Phosphorylated PD-1 participates in regulating CD28/Lck condensation.**

**A**: Condensates formed by pCD28-ICD/Lck_UD-SH3-SH2_ (AF488 labeled) with graded pPD-1-ICD (AF647 labeled) titration. Scale bar: 10 μm.

**B**: Coomassie blue staining and western blot verification of PD-1-ICDs with defined phosphorylated states. The localization of ITIM and ITSM sites on PD-1-ICD are shown.

**C**: Condensates formed by pCD28-ICD and Lck_UD-SH3-SH2_ in the presence of indicated PD-1-ICD variants. Scale bar: 20 μm.

**D**: Condensates formed by Lck_UD-SH3-SH2_ (AF488 labeled) with wild type (upper) and phosphorylated (lower) CD28-ICD on SLB after addition of ICDs of indicated co-inhibitory molecules. Scale bar: 10 μm.

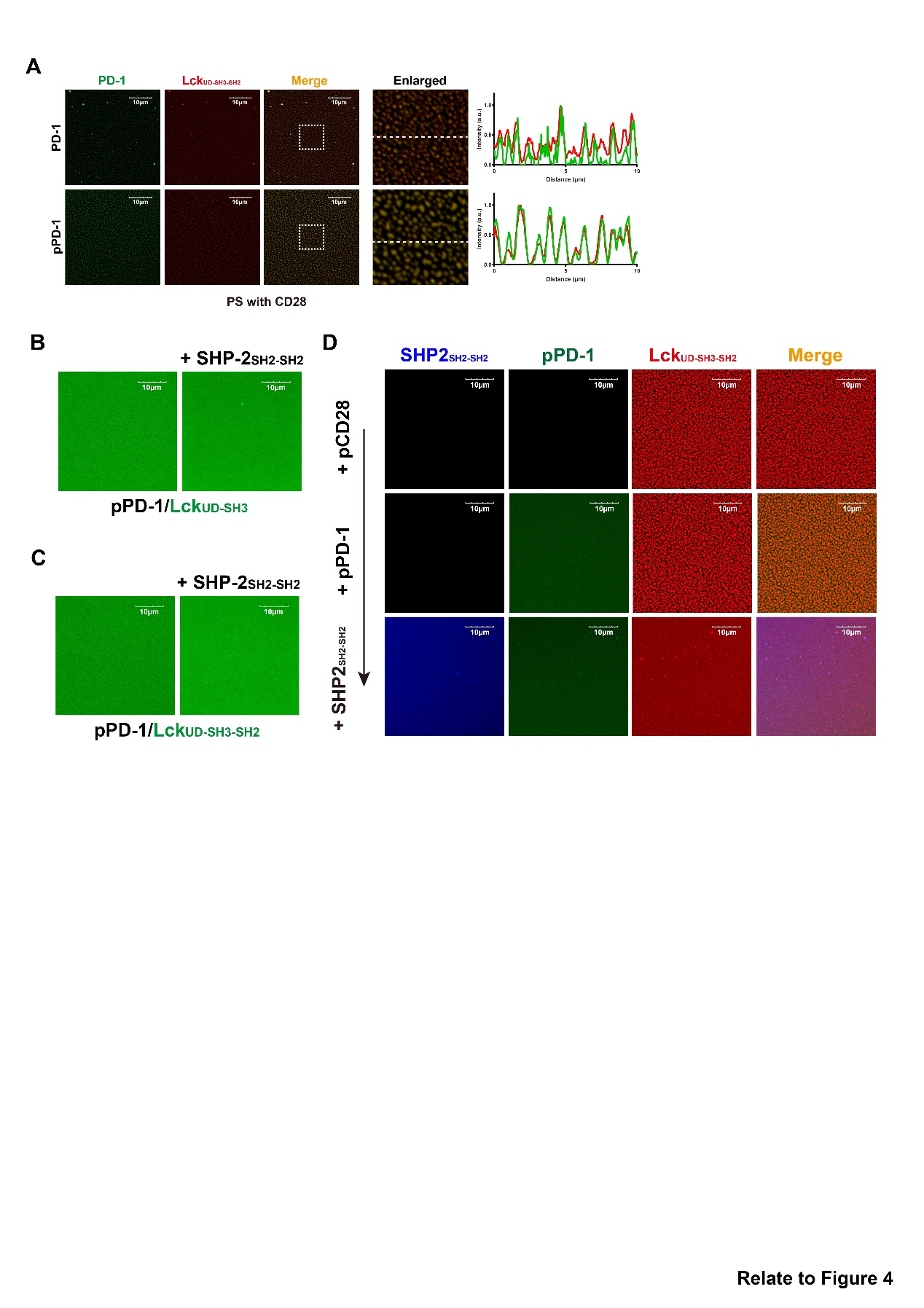

**Figure S6. SHP-2 facilitates pPD-1 to inhibit CD28/Lck phase separation.**

**A:** Localization of PD-1-ICD (upper) or pPD-1-ICD (lower) on SLB upon the Lck_UD-SH3-SH2_ clustering triggered by CD28-ICD addition. The fluorescence intensity profiles of PD-1-ICD (AF488 labeled) and Lck_UD-SH3-SH2_ (AF647 labeled) along the white dashed lines are plotted. Scale bar: 10 μm.

**B-C:** Lck_UD-SH3_ (**B**) and Lck_UD-SH3-SH2_ (**C**) clustering on SLBs in the presence of pPD-1-ICD and SHP-2_SH2-SH2_. Scale bar: 10 μm.

**D:** Condensates of pCD28-ICD/Lck_UD-SH3-SH2_ after sequential addition of pPD-1-ICD (AF488 labeled) and SHP-2_SH2-SH2_ (AF647 labeled). Scale bar: 10 μm.

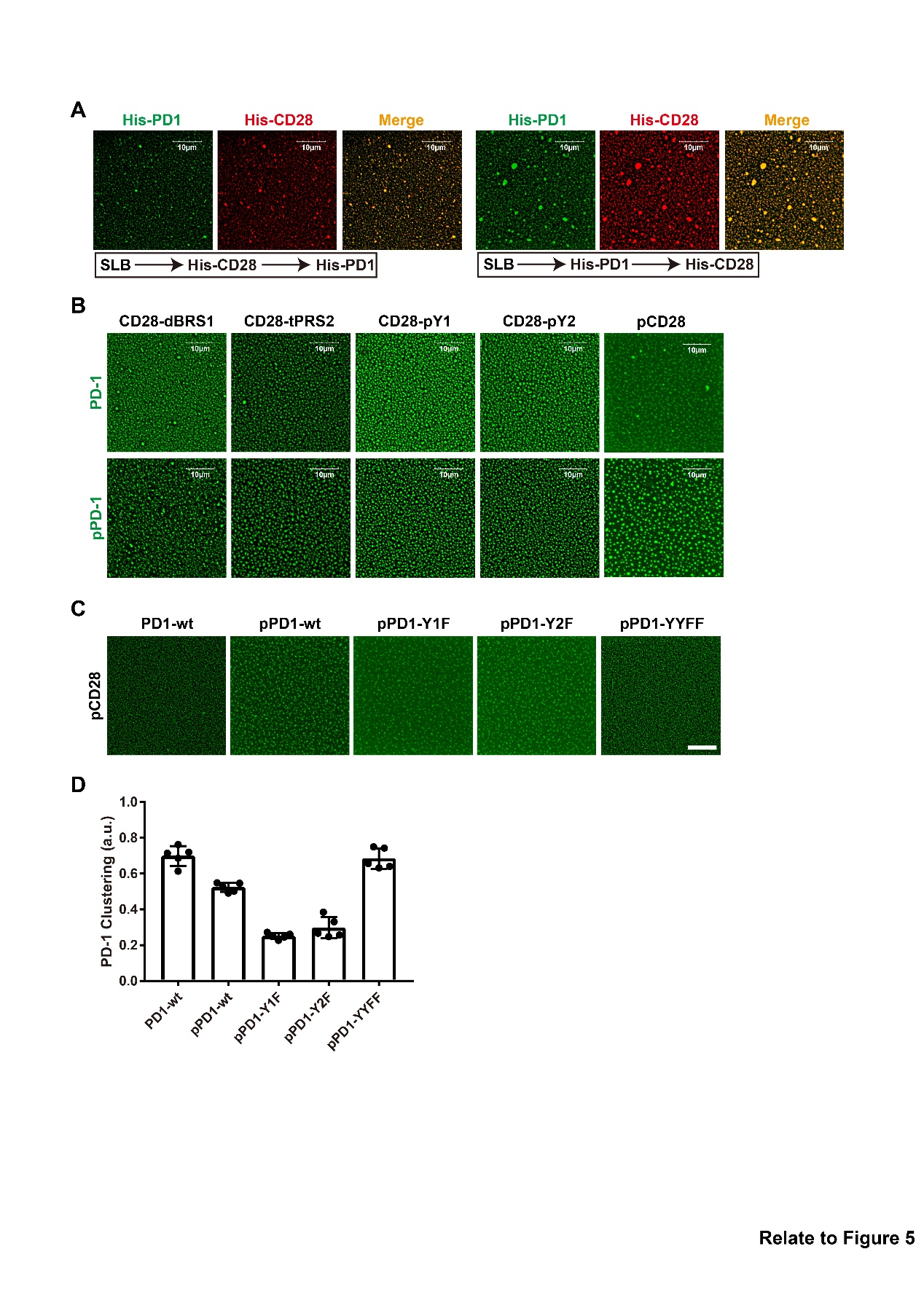

**Figure S7. Molecular mechanisms of CD28/PD-1 phase separation.**

**A:** Condensates formed by CD28-ICD (AF647 labeled) and PD-1-ICD (AF488 labeled) on SLBs with protein addition sequence shown.

**B:** Phase separation of PD-1-ICD (upper) or pPD-1-ICD (lower) with mutated or phosphorylated CD28-ICDs. Scale bar: 10 μm.

**C-D:** Representative images (**C**) and quantification (**D**) for pPD-1-ICD clustering induced by pCD28-ICD. Scale bar: 10 μm. Data are presented as Mean±SD, n=5.

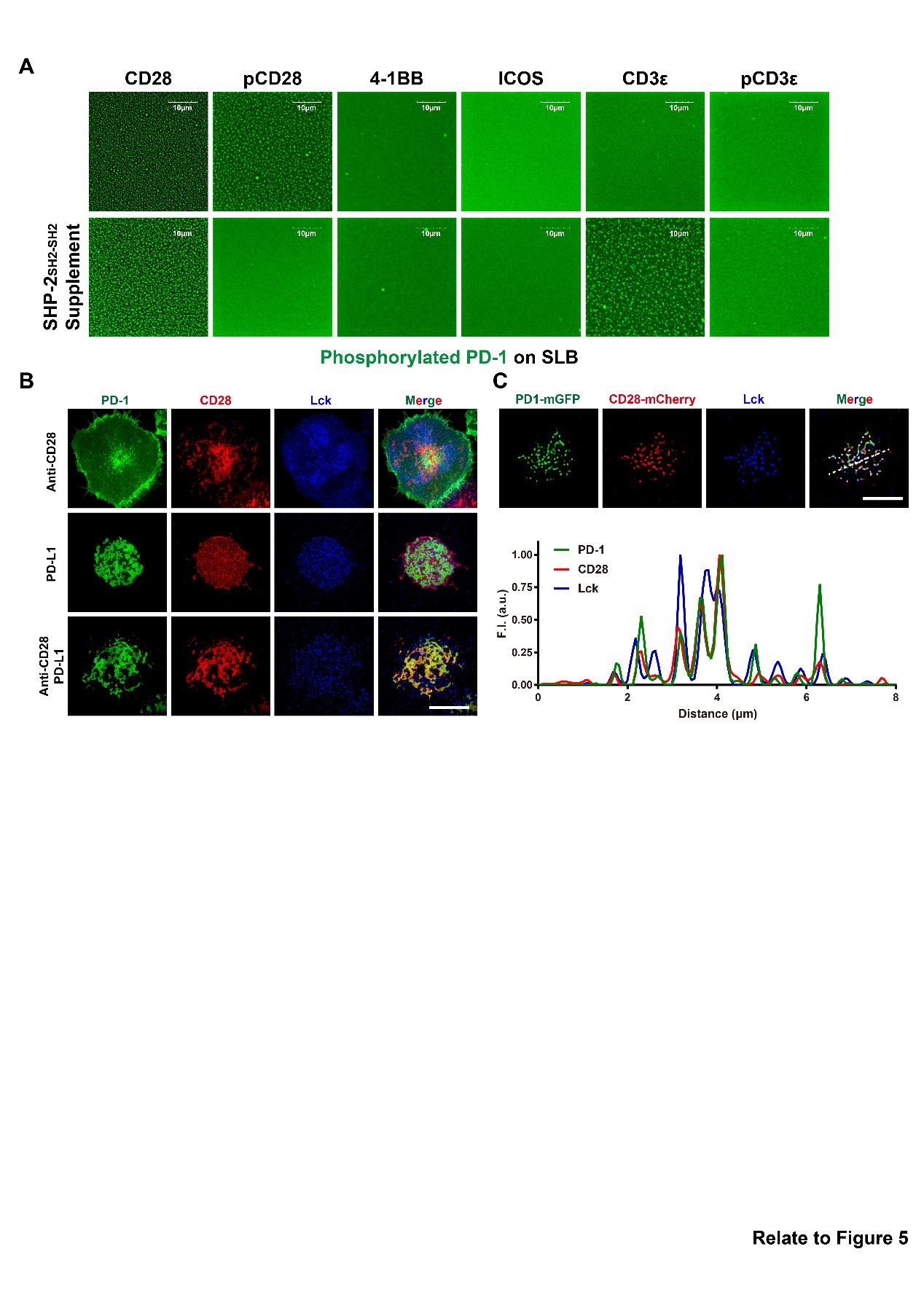

**Figure S8. PD-1 selectively phase-separates with CD28 and forms heterogenous condensates with CD28 and Lck.**

**A:** Clustering of pPD-1-ICD (AF488 labeled) with the indicated ICDs in the absence or presence of SHP2_SH2-SH2_ on SLBs. Scale bar: 10 μm.

**B-C:** Contact regions of Jurkat-CD28^mCherry^/PD-1^mGFP^ cells with SLBs (**B**) or glass surfaces (**C**) coated with anti-CD28 antibody, PD-L1 or both, cells are fixed and then stained with anti-Lck (APC labeled). The intensity of PD-1, CD28 and Lck are plotted alone the white dashed line in **C**. Scale bar: 10 μm (**B**) and 5 μm (**C**).

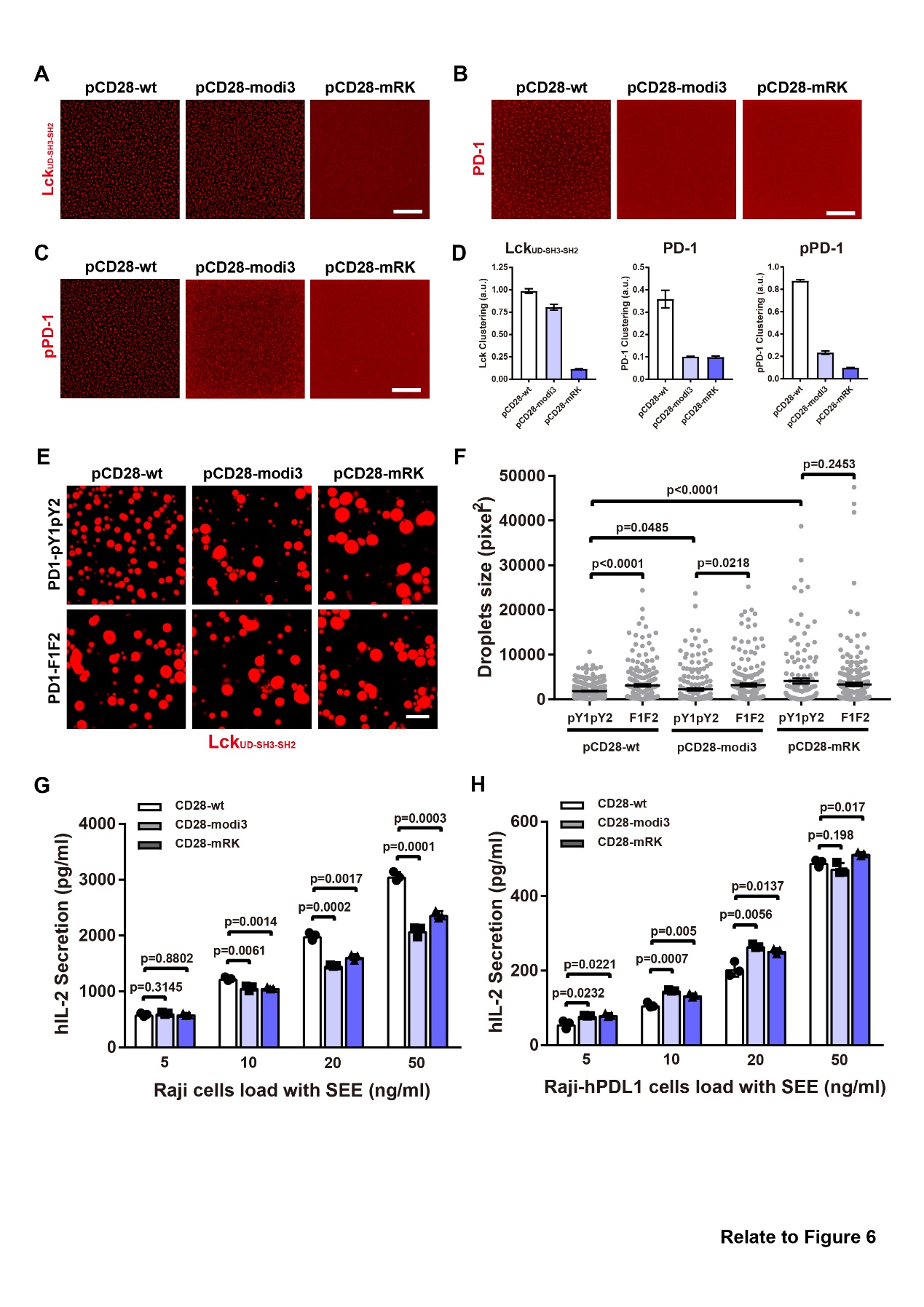

**Figure S9. CD28 mutations impair CD28/PD-1 phase separation and enhance T cell activation.**

**A-D:** Representative images (**A-C**) and quantification (**D**) of Lck_UD-SH3-SH2_ (**A**; **D**, left), PD-1-ICD (**B**; **D**, middle) and pPD-1-ICD (**C**; **D**, right) clustering on SLB induced by wild type or mutant CD28-ICDs. Scale bar: 10 μm. Data are presented as Mean±SD, n=5.

**E-F:** Representative images (**E**) and quantification (**F**) of pCD28-ICD/Lck_UD-SH3-SH2_ droplet size in the presence of the indicated pPD-1-ICDs. Scale bar: 20 μm. Data are presented as Mean±SEM.

**G-H**: IL-2 secretion of wild type or mutant Jurkat-CD28^mCherry^/PD-1^mGFP^ cells stimulated with SEE-loaded PD-L1 negative (**G**) or PD-L1 positive (**H**) Raji cells. Data are presented as Mean±SEM, n=3.

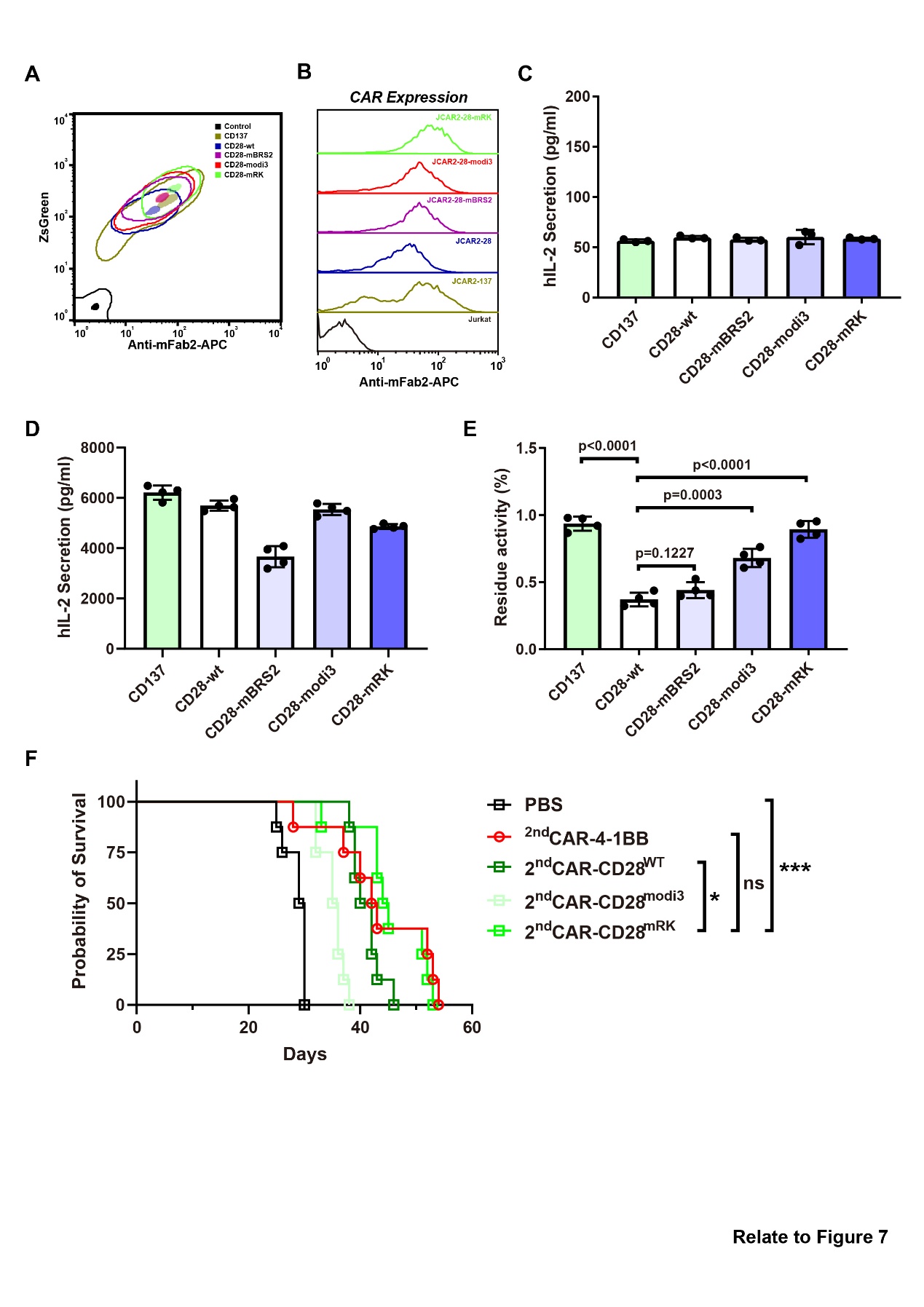

**Figure S10. CD28 mutants incorporated CAR-T cells improve safety and efficacy.**

**A-B:** Surface CAR expression on transduced Jurkat cells.

**C-E:** IL-2 secretion by second-generation CAR-Jurkat cells in the absence (**C**, n=3) or presence (**D**, n=4) of Raji cells. Quantification of residue activity upon PD-1 triggering are shown (**E**, n=4). The raw IL-2 secretion of second-generation CAR-Jurkat cells stimulated with PD-L1 negative (IL2) or positive (IL2^PD-L1^) Raji cells are showing in (**D**) and Figure 7C, the residue activity is calculated as (IL2- IL2^PD-L1^)/IL2. Data are presented as Mean±SD.

**F:** Survival probability of NSG mice treated with indicated CAR-T cells (n=8).
